# Metabolic consequences of various fruit-based diets in a generalist insect species

**DOI:** 10.1101/2022.10.21.513142

**Authors:** Laure Olazcuaga, Raymonde Baltenweck, Nicolas Leménager, Alessandra Maia-Grondard, Patricia Claudel, Philippe Hugueney, Julien Foucaud

**Affiliations:** UMR CBGP (INRAE-IRD-CIRAD, Montpellier SupAgro), Campus International de Baillarguet, CS 30 016, 34988 Montferrier / Lez cedex, France; Department of Agricultural Biology, Colorado State University, Fort Collins, Colorado 80523, USA; Université de Strasbourg, INRAE, SVQV, UMR A 1131, F-68000 Colmar, France

**Author notes:** These authors contributed equally to this work (Laure Olazcuaga, Raymonde Baltenweck, Philippe Hugueney, Julien Foucaud).

**Keywords:** niche breadth, ecological specialization, metabolomics, generalism, fruit diet, host plant, spotted wing drosophila, Drosophila suzukii

## Abstract

Most phytophagous insect species exhibit a limited diet breadth and specialize on few or a single host plant. In contrast, some species display a remarkably large diet breadth, with host plants spanning several families and many species. It is unclear, however, whether this phylogenetic generalism is supported by a generic metabolic use of common host chemical compounds (‘metabolic generalism’) or alternatively by distinct uses of diet-specific compounds (‘multi-host metabolic specialism’)? Here, we simultaneously investigated the metabolomes of fruit diets and of individuals of a generalist phytophagous species, *Drosophila suzukii*, that developed on them. The direct comparison of metabolomes of diets and consumers enabled us to disentangle the metabolic fate of common and rarer dietary compounds. We showed that the consumption of biochemically dissimilar diets resulted in a canalized, generic response from generalist individuals, consistent with the metabolic generalism hypothesis. We also showed that many diet-specific metabolites, such as related to the particular color, odor or taste of diets, were not metabolized, and rather accumulated in consumer individuals, even when probably detrimental to fitness. As a result, while individuals were mostly similar across diets, the detection of their particular diet was straightforward. Our study thus supports the view that dietary generalism may emerge from a passive, opportunistic use of various resources, contrary to more widespread views of an active role of adaptation in this process. Such a passive stance towards dietary chemicals, probably costly in the short term, might favor the later evolution of new diet specializations.

## Introduction

One of the most salient features of living organisms is their diversity in term of ecological requirements. Some species show extreme specialization to their environment (e.g., obligate symbionts), while others are able to use a wide range of resources and tolerate many environmental conditions. Furthermore, these relationships between organisms and their biotic and abiotic environments are dynamic at various spatial and temporal scales. As such, the concepts of niche breadth and ecological specialization are of paramount importance in ecology and evolutionary biology (Futuyma & Moreno, 1988; Sexton et al., 2017).

The dietary requirements of phytophagous insects are a textbook example of variance in niche breadth (Forister et al., 2012; Futuyma & Moreno, 1988; Jaenike, 1990). In this group, the vast majority of species are specialists of one family of plants (Forister et al., 2015) and this specialization is often accompanied by extreme morphological and/or physiological specializations to exploit a particular plant (e.g., Berenbaum, 1981; Hotti & Rischer, 2017). In contrast, some rarer species display a much wider niche breadth, allowing them to consume up to several dozen plant families (Clarke et al., 2005; Forister et al., 2015). The larger proportion of specialist than generalist species (Forister et al., 2015), as well as the fact that transitions between specialism and generalism are bidirectional (Day et al., 2016; Janz et al., 2001; Nosil, 2002; Nylin et al., 2014), raises questions about the origin and maintenance of generalism (Forister et al., 2012; Singer, 2008), and its biochemical basis.

From a physiological point of view, dietary generalism has been much less experimentally studied than specialism, probably in part because it requires the simultaneous consideration of numerous host species and the tracking of physiological responses that may involve far more components than specialism. Additionally, study of dietary generalism lacks the intellectual appeal of bi-species interactions leading to biochemical and genetic arm-races. Indeed, many research scrutinized the evolution of plant defenses and their corresponding detoxifying pathways in herbivores (e.g., the glucosinolate–myrosinase system of Brassicaceae countered by *Spodoptera frugiperda*, Winde & Wittstock, 2011). Comparatively, knowledge about how generalist species cope with a wide variety of plant biochemistry is scarce (but see Roy et al., 2016).

The core question of dietary generalism at the level of the organism is how a generalist species metabolizes different diets, which involves some basic issues that are still pending. For instance, do individuals of a generalist species have a unique biochemical response to all diets or do they differ according to their diet? In evolutionary terms, do dietary generalist species exhibit a canalized response to different dietary environments through a single, neutral and generic use of hosts (H1; hereafter termed ‘*metabolic generalism*’), or do they show multiple specialized, adapted uses of different hosts (H2; hereafter termed ‘*multi-host metabolic specialism*’)? Broad organismal-level patterns of biochemical composition can help answer this question because individuals on different diet should be chemically similar across diets according to the former hypothesis, or mostly dissimilar according to the latter (Figure 1).

**Figure 1:**
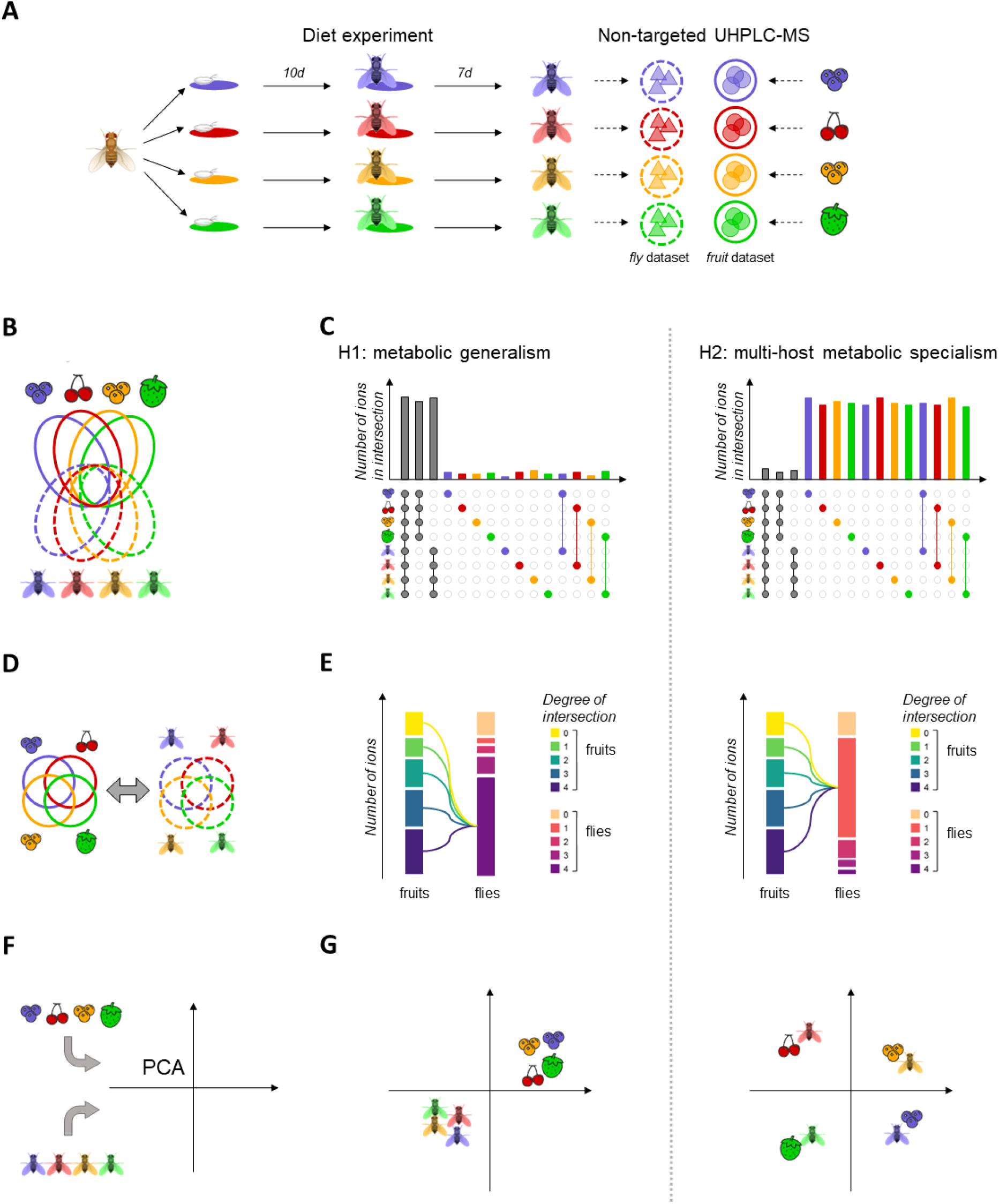
Schematic overview of experimental design, host use analyses and expectations according to the ‘*metabolic generalism*’ and ‘*multi-host metabolic specialism*’ hypotheses. In order to investigate whether different food sources were treated similarly or differently by a generalist species, we followed a simple experimental procedure (A), where eggs of a single founder population were separately developed on four fruit purees for 10 days and kept on the same fruit for seven days of adult life. Both adult flies and fruit purees were processed to obtain the *fly* and *fruit* dataset, with respective ions represented by colored triangle and dots. These datasets were used to perform three analyses (B, D, F) linked to three diverging expectations under our competing hypotheses of ‘*metabolic generalism*’ and ‘*multi-host metabolic specialism*’ (C, E, G). First, the comparison of the relative sizes of intersections of the complete dataset (B) should indicate whether ions are mostly shared between intersections of large sizes such as all fruits, all flies or all samples (C, left) or whether ions are mostly confined to intersections of small size such as discrete categories or fruit-fly pairs (C, right). Second, relationships between intersections between fruits on one hand and between flies on the other (D) should indicate whether common or rare fruits ions are commonly (E, left) or rarely (E, right) found in flies. Third, a PCA of the complete dataset should indicate whether fruits and flies form distinct metabolic clusters (F, left) or whether fruit-flies pairs cluster away from each other (F, right). Regarding expectations (C, E, G), all left and right panels are consistent with the ‘*metabolic generalism*’ and ‘*multi-host metabolic specialism*’ hypotheses, respectively.

To refine our understanding of diet generalism, we here use an untargeted metabolomics approach to characterize the metabolic impact of various fruit-based diets on the generalist insect species *Drosophila suzukii* (Figure 1A). The range of molecular compounds detected and quantified by current metabolomic methods may span many hundreds or thousands of molecules, ensuring a high-quality snapshot of the individuals’ physiological state (Johnson et al., 2016). Additionally, untargeted metabolomics provides the opportunity to directly compare diets and individuals that consume them, in an objective and repeatable way. The invasive *Drosophila suzukii* (the spotted wing drosophila) displays an exceptional diet breadth of over 20 plant families (Asplen et al., 2015; Bellamy et al., 2013; Cini et al., 2014; Kanzawa, 1939; Kenis et al., 2016; Lee et al., 2011, 2015; Poyet et al., 2015). While multiple experimental studies have found that different diets have an influence on its larval development (Kenis et al., 2016; Olazcuaga et al., 2019; Poyet et al., 2015), it is widely accepted that *D. suzukii* is a diet generalist species.

By directly comparing the metabolomes of fruits and flies that consume them, we investigated the following questions: (1) How biochemically different are the selected major host plant species used by *D. suzukii*? (2) Do generalist individuals use different host plants in a similar way, qualitatively and quantitatively? (3) Can we discriminate individuals of a generalist species according to their diet, and identify diet-specific metabolites in generalist individuals?

After having confirmed that the investigated host plants exhibit large biochemical differences, metabolomes of flies that experienced specific diets during their larval and early adult stages were compared qualitatively and quantitatively (Figure 1). This comparison of various sets of fruit and fly metabolites confirmed that diet generalism is mainly due to a generic, diet-unspecific qualitative response to host biochemistry, i.e., ‘metabolic generalism’. This tolerant reaction to diet entailed the passive accumulation of distinctive fruit metabolites in flies, that could easily be assigned to their diet.

## Results

The metabolomes of four major crop fruits (blackcurrant, cherry, cranberry and strawberry) and their corresponding flies (that consumed them both at the larval and early adult stages) were characterized simultaneously using ultra-high-performance liquid chromatography coupled to mass spectrometry (UHPLC-MS; Figure 1A). Our untargeted approach enabled the detection and relative quantification of a set of 11,470 unique ions that were found in at least one fruit or fly sample. While the simultaneous processing of fruit and flies’ samples provided a single *integral* dataset enabling direct comparison between diets and consumers, some analyses required the fruit-related and fly-related data to be separated, resulting in a *fruit* dataset and a *fly* dataset. Although the *integral*, *fruit* and *fly* datasets are based on ions characterized by UHPLC-MS, some ions will be referred to as “compounds” or “metabolites” later in the manuscript, depending on the context.

### Metabolomic analysis of fruit composition

A preliminary requirement to investigate dietary generalism is to directly compare diets from a biochemical perspective, because different plant families may share chemical properties that could aid transition from specialism to generalism (e.g., share the same plant defense compounds; Bowers, 1983). To exclude such misleading generalism, an often overlooked question is therefore whether the taxonomic scale (e.g., species) fairly represent different dietary challenges. We evaluated the biochemical proximity of the four investigated fruits using a Principal Component Analysis (PCA) framework on the *fruit* dataset. This analysis showed that the four investigated fruits were dissimilar and that technical replicates were highly similar (Figure 1—figure supplement 1). Blackcurrant was very dissimilar to all other fruit (discriminated on the first dimension of the PCA explaining 35% of the *fruit* dataset variance). Cherry was then discriminated by the second dimension of the PCA (28% variance explained), and cranberry and strawberry by the third dimension of the PCA (25% variance explained). Furthermore, qualitative data alone enabled perfect discrimination between fruit, with several hundred ions diagnostic of each investigated fruit (Figure 1—figure supplement 2). The distinctive biochemical signatures of the four investigated fruits thus provide ideal ground to decipher the effect of their use on their generalist consumers.

Additionally, we checked whether fruit-derived ions detected in flies were related to their development on a specific diet (i.e., the metabolization of fruit) or, conversely, may come from direct contamination of the flies by fruits. The absence of direct contamination was confirmed by the absence of correlation between ion quantities in fruits and flies that consumed them (all Kendall’s τ < 0.2; Figure 1—figure supplement 3).

### Host use

Whether individuals of a generalist species process all types of diet in a similar, generic way (‘metabolic generalism’) or show specific pathways to metabolize unique compounds of different diets (‘multi-host metabolic specialism’) entails largely non-overlapping expectations for both qualitative and quantitative biochemical composition of individuals relatively to their diets (Figure 1B & C).

A first approach lies on the fact that metabolic generalism is expected to result in particular patterns of overlap between ions, when all fruits and flies are considered together (Figure 1B, Analysis 1). If *D. suzukii* is a metabolic generalist, flies should all share a large part of their ions, fly ions specific of a particular diet should be rare or absent, and/or ions specific of single fruit-fly pair (i.e., a pair of fruit and flies grown over the same fruit) should also be rare (Figure 1C Analysis 1, left panel). On the contrary, multi-host metabolic specialism would be detected by any given fruit-fly pair sharing large amounts of ions but not with other pairs, and/or large amounts of ions found only in flies that are specific of each diet (Figure 1C Analysis 1, right panel). To test these predictions, we computed intersection sizes between all fruits’ and flies’ ions from a presence/absence (qualitative) dataset derived from our quantitative *integral* dataset. Comparison of the overall intersections of ions present in the different fruits and the flies grown over these fruits showed that most ions are shared between large groups of fruits and flies (*i.e.,* with higher degrees of intersection; Figure 2A). The largest intersection includes 3754 ions shared by all fruits and all flies (33%). The second largest intersection comprises 1030 ions shared between all flies (and absent in fruits; 9%). The third largest intersection contains 463 ions shared between all fruits (and no flies; 4%). High-order degree characterizes all other largest intersections, except for a set of 297 ions uniquely found in blackcurrant fruits. Figure 2B focuses on the sets of ions found uniquely in fruits, flies or fruit-fly pairs (i.e., the intersections of degree 1 and 2). While ions found uniquely in each fruit are not uncommon (>100 ions for all fruits), ions found uniquely in flies raised on a particular diet are rare (9 in cherry-fed flies, 6 in cranberry-fed flies, 1 in blackcurrant-fed flies and none in strawberry-fed flies). Ions that are distinctive of a particular fruit-fly pair are also seldom, ranging from 77 in the blackcurrant fruit-fly pair to only 12 in the strawberry fruit-fly pair.

**Figure 2:**
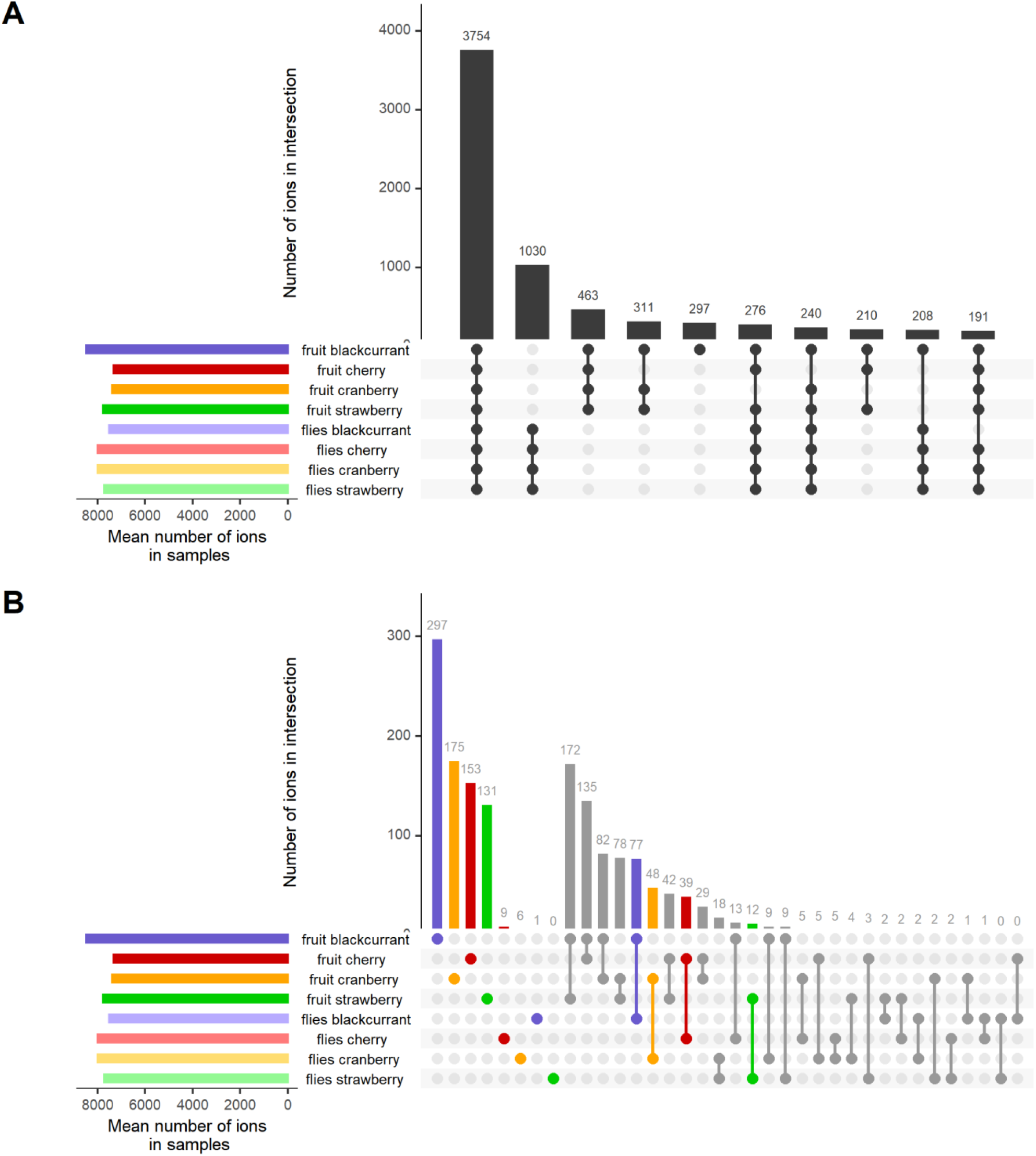
Global trends of shared ions between all fruits and all flies. (A) the ten largest intersections between all metabolomes comprise many high degree intersections, including ions shared between all fruit and all flies’ samples (#1), ions, shared between all flies (#2) and ions shared between all fruits (#3). A notable exception to this pattern is the presence of ions found only in the blackcurrant fruits (#5). (B) A focus on ions unique to particular samples or sample pairs (i.e., intersections of degree 1 and 2, respectively) illustrate that, while ions unique to fruits are found in high numbers, ions unique to diet in flies are largely absent. Ions specific to a given fruit-flies pair are infrequent. See Figure 1C for expectations according to diet generalization hypotheses.

Similarity and dissimilarity in the use of different host plants could stem from the metabolic fate of common and unique fruit metabolites when ingested by flies. Differently-fed flies metabolizing compounds that are shared between fruits to produce metabolites shared between flies would be indicative of ‘metabolic generalism’ (Figure 1C, Analysis 2). On the opposite, differently-fed flies metabolizing compounds private to a given fruit to produce new metabolites (mostly not shared between flies) would be indicative of ‘multi-host metabolic specialism’. We thus inspected of the intersection degrees of all fruits and the intersection degrees of all flies, together with their relationships (Figure 3). First, the intersection of all fruits (left column) is largely formed from ions common to all fruits (48.6% of all ions present in fruits). Ions present in lesser degree intersections are found in proportions roughly equal to each other (20.1%, 14.8% and 16.5% for intersections of degree 3, 2 and 1, respectively). Approximately 10% of all ions found in this study are not detected in fruits, but only in flies (yellow stack, left column). Second, the intersection degrees of flies show marked differences with that of fruits (Figure 3, right column). These include a higher proportion of metabolites common to all flies (74.7% of all ions present in flies), smaller proportions of lesser degree intersections (intersections of degree 3, 2 and 1 represent 7.8%, 6.1% and 11.3% of all ions present in flies, respectively), and a large proportion of fruit metabolites that are absent in flies (21.2% of all detected ions). Third, the inspection of the relationships between fruit and flies’ metabolomes shows that most common fruit ions are the largest, but not unique, part of the most common fly ions. It is important to note that 86% of ions that were present in flies but not in any fruit (i.e., derived from *de novo* produced metabolites; yellow stack, left column) are ions common to all flies regardless of their diet (Figure 3, yellow flow from 0 to 4). In contrast to the most common fruit ions, ions unique to a single fruit or shared by a few fruits (fruit intersection degrees 1 to 3) roughly follow the same distribution of 1/3 being shared by all flies, 1/3 being shared by a few flies, and 1/3 being shared by no flies. This includes ions unique to a given fruit, 35.2% of which are found in all flies. Fly ions that are specific of a given diet come from fruit ions with various intersection degrees. In particular, 25.7% are common to all fruits, 22.6% are unique to a single fruit and only 1.6% are produced *de novo* (16 ions).

**Figure 3:**
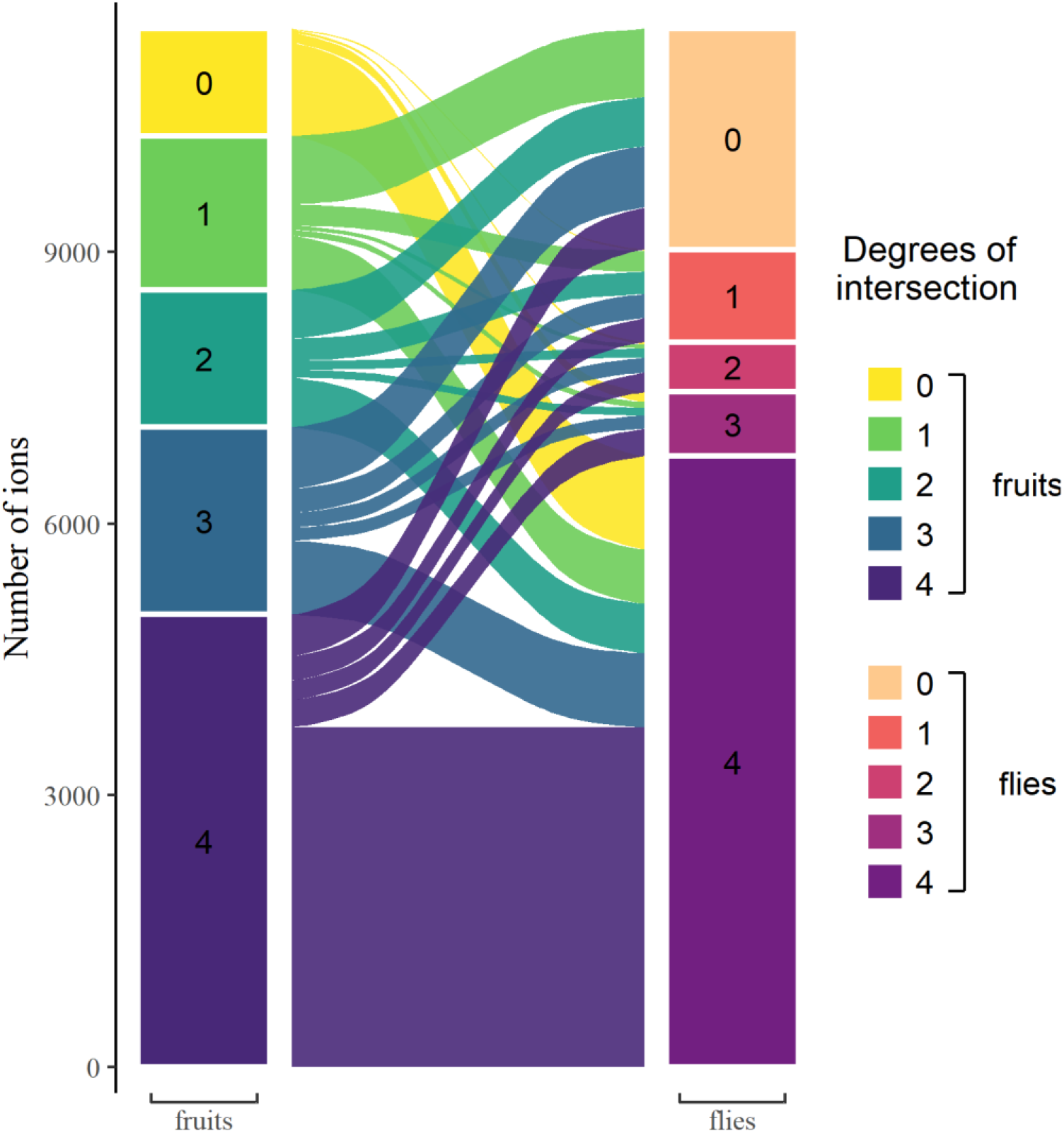
Relationships between ions respectively found in fruits and flies, according to their degree of intersection. Ions that are shared within fruits or within flies show a higher degree of intersection. Most ions found in flies are found in all flies, but not necessarily in all fruits. It is also noteworthy that almost all ions produced *de novo* by flies are found in all flies, regardless of their diet. See Figure 1E for expectations according to diet generalization hypotheses.

Finally, we visually investigated the multi-dimensional clustering of fruits’ and flies’ metabolomes using a PCA using the *integral* dataset (Figure 1B, Analysis 4). If diets are the main influence on flies’ metabolomes, flies should not cluster together and rather localize close to their corresponding fruit (Figure 1C, Analysis 4). On contrary, if flies similarly process different diets, flies’ metabolomes should cluster together, away from the fruits. Results of the PCA show that while different fruits are still easily discriminated, differently-fed flies all clustered into a single group (Figure 3—figure supplement 1).

Altogether, our analyses underscore that the metabolomes of flies fed on different fruits are mostly identical, especially from a qualitative standpoint, and unanimously point to metabolic generalism, over multi-host metabolic specialism, as a mechanism promoting diet breadth.

### Metabolomic discrimination of diets in flies

Given the overall qualitative similarity of flies that developed on different diets, we then investigated whether quantitative hallmarks of diet consumption could be detected in flies. A PCA focusing on the *fly* dataset demonstrated that flies’ diet can be readily inferred (Figure 4). While the first dimension of the PCA (21.6% of variance explained) discriminates between biological replicates of our experiment (i.e., generations of flies), the second and third dimensions of the PCA (17.5% and 9.4% of variance explained, respectively) discriminate fly metabolomes according to their diets.

**Figure 4:**
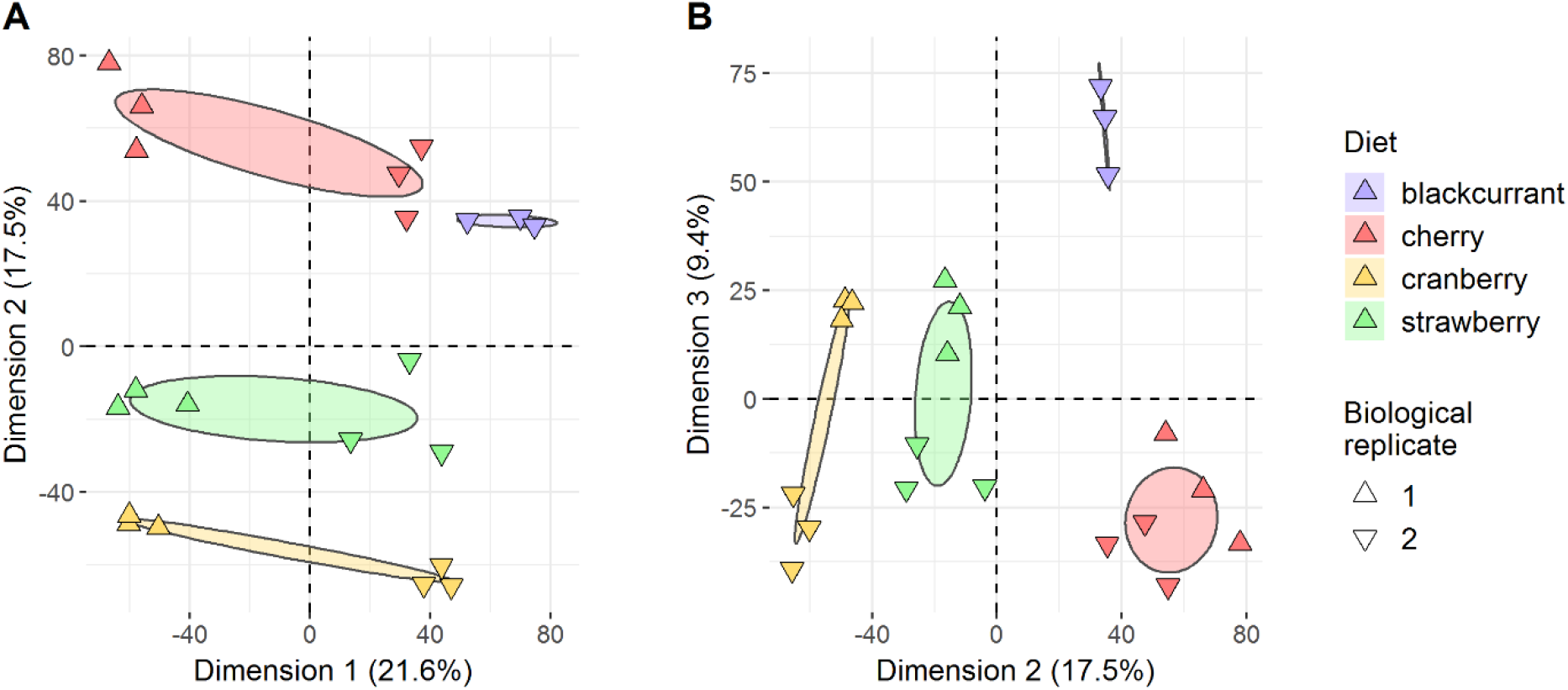
Flies’ metabolomes differ according to diet. Flies’ samples only were included in this PCA. Differently oriented triangles indicate biological replicates (fly generations). (A) Dimensions 1 and 2; (B) Dimensions 2 and 3.

To further detect which ions were indicative of the development on particular fruits, we then computed diet-specific binomial linear models with Elastic Net regularization. We specifically aimed at producing two lists of diet-specific ions based on their quantitative levels in flies’ metabolomes and on two sets of parameters designed (*i*) to optimally classify flies according to their diet (‘large’ list) and (*ii*) to be as short as possible to allow manual curation of the identified ions (‘compact’ list, see Materials & Methods for details). This statistical approach yielded a ‘large’ list consisting of 162 fly ions associated with a blackcurrant diet, 98 with a cherry diet, 111 with a cranberry diet and 37 with a strawberry diet. All four ‘large’ models enabled a perfect classification of flies according to their diet (ratio of deviance explained >98%). As expected, these ions are part the top set of ions driving the explained variance in the dimensions of the previous PCA, as shown by their location in the ion space (Figure 4—figure supplement 1). The ‘compact’ set of Elastic Net parameters attained similar deviance ratio and yielded a list of 64 fly ions associated with a blackcurrant diet, 38 with a cherry die t, 46 with a cranberry diet and 12 with a strawberry diet. All fly ions in the ‘compact’ list were present in the ‘large’ list. As shown in Figure 4—figure supplement 2, the ‘large’ list of fly ions can perfectly class fly samples according to their diet. It also suggests that most of these fly ions are present in similar quantities in their respective fruit, with few exceptions. Consistent with these results, we verified that ions of the ‘large’ list for a given flies’ diet were significantly more present in the flies of the said diet than in the flies developed on other diets and also more present in the said fruit than on other fruits (Figure 4—figure supplement 3). Depending on the diet, quantities of these distinctive ions found in flies could be correlated to the quantity found in the fruit (*e.g.,* blackcurrant, cherry; Figure 4—figure supplement 4A & B) or not (*e.g.,* cranberry, strawberry; Figure 4—figure supplement 4C & D).

Diet-associated ions were found to be part of intersections of every degree, both in flies and in fruits (Figure 4—figure supplement 5). In flies, large parts of these ions were specific of the flies that consumed the diet, especially for blackcurrant-fed flies (degree 1: 70%, 45%, 41% and 35% in blackcurrant, cherry, cranberry and strawberry-fed flies, respectively). However, ions contained in intersections of larger degrees were also observed, including ions found in every fly (degree 4: 14%, 34%, 32% and 30% in blackcurrant, cherry, cranberry and strawberry-fed flies, respectively; Figure 4—figure supplement 5 & 6). While these diet-distinctive metabolites did not always originate from fruit-specific metabolites and could be detected in other flies and other fruits (Figure 4—figure supplement 5), their quantities were lower there than in the focal fruit and flies who consumed it. It is worth noting that few diet-specific ions were produced *de novo* by flies (fruits intersection of degree zero: 0%, 2%, 4% and 0% ions in blackcurrant, cherry, cranberry and strawberry-fed flies, respectively), consistent with the metabolic generalism hypothesis.

Altogether, we found that quantitative levels of ions are critical to correctly classify fly metabolomes according to diet. Consistent with this finding, we observed that qualitative and quantitative datasets did not have the same resolving power when it came to distinguish flies based on their diet (Figure 4—figure supplement 7). While randomly picking ∼300 ions would lead to a 75% classification accuracy for flies’ diet when using their quantification levels, a similar performance was attained with a sampling of ∼1,000 ions when using only their presence. Regardless, flies’ diet could be readily inferred based on a relatively small subset of ions.

### Identification and fate of metabolites associated to fruit diets

To gain insight about distinctive features of flies and their diets, we tried to identify diet-specific fly ions of the ‘compact’ list together with the 50 major ions (which display the largest peak area) for each investigated fruit. Manual curation and detailed mass spectra analysis allowed us to identify 71 metabolites, 26 of which were confirmed with commercial standards. Identified metabolites belonged to major plant metabolite families, including anthocyanins, flavonoids, organic and hydroxycinnamic acids and terpenes (Supplementary Table 1). A two-dimensional clustering analysis of metabolites associated to diet illustrates that even a limited set of these compounds was sufficient to infer flies’ diet (Figure 5). A survey of the published literature showed that many of the metabolites identified here in specific fruits (and flies that were grown on them) had already been detected and identified in the same fruits (Figure 5 & Supplementary Table 1), thereby supporting their correct identification.

**Figure 5:**
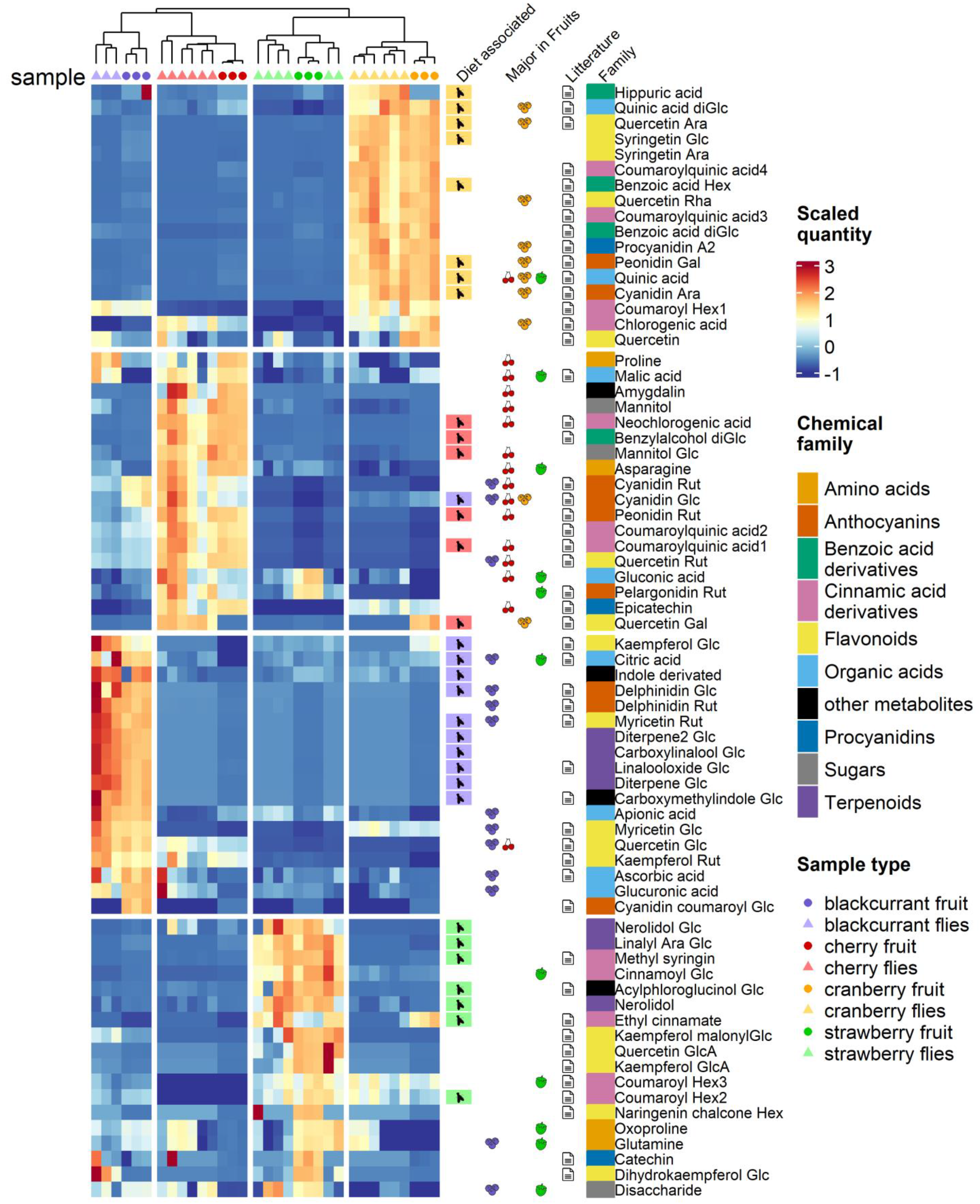
Heatmap of the relative quantity of identified metabolites in each fruit and fly sample. Heatmap scale colors indicate the relative quantity of each metabolite (row). Additional columns represent diet specificity in flies, major presence (top 50) in fruits, presence in literature in relation to one or several of the investigated fruits, and chemical family of each identified metabolite (see Supplementary Table 1 for detailed bibliographic information).

Depending on fruits and specific metabolites, relative amounts of fruit-derived metabolites detected in flies did not necessarily mirror their relative abundance in fruits (Figure 5—figure supplement 1). Indeed, some major fruit metabolites, such as the anthocyanins delphinidin in blackcurrant, cyanidin in cherry or pelargonidin in strawberry, were detected in flies, albeit with relatively low relative abundance (fly/fruit ratio between 0.01 and 0.001). Conversely, other less abundant fruit metabolites, such as terpenes, were relatively abundant in flies (fly/fruit ratio > 0.5). Despite the relatively low number of identified metabolites, our quantification results demonstrate that only some fruit metabolites are metabolized (or excreted), while others tend to accumulate in flies.

As expected from previous studies, organic acids tended to be shared by all fruits, but with a particular balance depending on fruits: malic, quinic and citric acids were always present in fruits (and flies) but with elevated levels in cherry, cranberry and blackcurrant fruits (and flies that consumed them), respectively. Another chemical family ubiquitous in plants is flavonoids, with many potential physiological effects for consumer organisms, such as neurobehavioral modulation of activity (Bugel & Tanguay, 2018), feeding and oviposition (Simmonds, 2001; Treutter, 2006). All fruits harbored major flavonoids such as quercetin, kaempferol or myricetin, with specific substituent groups: quercetin arabinoside and rhamnoside in cranberry, quercetin rutinoside in cherry, quercetin glucoside in blackcurrant and quercetin glucuronide in strawberry. Those substituents have been shown to modulate the bioactivity of flavonoids (Bugel & Tanguay, 2018) and might mediate differential developmental responses to fruit diet in *D. suzukii* (Olazcuaga et al., 2019). While these substituents can undergo profound modifications by the gut microbiome (e.g., hydrolysis), we found no large-scale trace of microbiota influence in the present study, with flavonoids being essentially carried ‘as is’ in flies’ metabolomes. As an interesting subclass of flavonoids, major anthocyanins characteristic of the investigated fruits (and of their color) were readily found in flies that consumed these fruits. Strawberry-fed flies displayed elevated levels of pelargonidin glucoside, the main anthocyanin of strawberry fruits (da Silva et al., 2007). Similarly, cranberry- and cherry-fed flies presented high levels of cyanidin arabinoside, and cyanidin glucoside and rutinoside, the main anthocyanins found in cranberry and cherry, respectively (Acero et al., 2019; Seeram et al., 2004). Finally, fruits and flies that consumed them also displayed specific chemical signatures associated to terpenoids, considered as the largest class of plant secondary metabolites (Dudareva et al., 2004). Our results show that blackcurrant-fed flies can easily be discriminated by their levels in some linalool derivatives. While this volatile, floral-scented monoterpene is found in many plant families (Knudsen et al., 2006; Pichersky & Lewinsohn, 2011), specific linalool derivatives, e.g., carboxylinalool glycoside and linalooloxide glycoside, were predictive of the flies’ blackcurrant diet. Similarly, flies that developed on strawberry can be differentiated based on their elevated content in nerolidol, a sesquiterpene found in the floral scent of many plant families (Knudsen et al., 2006) and in fruits including strawberry (Aharoni et al., 2004).

While most of these signatures involve passive accumulation of plant secondary metabolites, we also uncovered traces of the metabolization of fruits compounds. Indeed, we found elevated levels of hippuric acid in all flies, and particularly in cranberry-fed flies. Hippuric acid originates from the metabolization of various phenolic compounds (such as flavonoids and chlorogenic acids) by the gut microbiome, and is a biomarker of fruits (and especially berries) consumption in various species including humans (De Simone et al., 2021; Ulaszewska et al., 2020). This result highlights that the metabolization process of major fruit compounds by *D. suzukii* may be relatively universal, and lack specificity, as shown with the accumulation of monoterpenes in blackberry-fed flies. However, it is noteworthy that some compounds belonging to other families of toxic plant metabolites do not seem to affect *D. suzukii*. Both cherry puree and cherry-fed flies were found to contain amygdalin, which belongs to the cyanogenic glucoside family. The toxicity of cyanogenic glucosides has been widely documented in the context of plant-insect interactions (Zagrobelny et al., 2004). However, cherry has been identified as an excellent host for *D. suzukii*, in both oviposition preference and larval performance experiments (Olazcuaga et al., 2019).

## Discussion

The direct comparison of metabolomes of fruits and of individuals that were grown on them sheds some light about basic issues pertaining to diet generalism (Forister et al., 2012).

The biochemical composition of stone fruits and berries has been explored in numerous studies, highlighting the major contributions of sugars, organic acids and secondary metabolites to the chemistry of fruits. Our untargeted overview of fruit composition enabled us to confirm that these fruits differ widely in their biochemical composition, and most importantly, that phylogenetic differences actually correspond to different biochemical landscapes for frugivorous species.

The use of diverse hosts had a variety of biochemical consequences on individuals of the generalist *D. suzukii*, all consistent with the ‘metabolic generalism’ hypothesis. Qualitatively, individuals showed a remarkably fixed chemical response to diet, coherent with a canalized response to this environmental parameter. This pattern mirrored the metabolic homeostasis found in generalists *Drosophila* flies reared on various protein:carbohydrates (P:C) ratios (Silva-Soares et al., 2017; Watanabe et al., 2019; Young et al., 2018). Quantitatively, however, different diets had a significant impact on the levels of a number of ions, resulting in the straightforward identification of flies’ diet based on their metabolome. Our study thus highlights that the physiological response to various diets, while generic in structure, was plastic when relative metabolite levels were considered. This biochemical response could be expected from studies underlining the transcriptional plasticity in generalist species from all major insect clades (Birnbaum & Abbot, 2020; Celorio-Mancera et al., 2016; Celorio-Mancera et al., 2012, 2013; Christodoulides et al., 2017; Ragland et al., 2015; Simon et al., 2015).

The simultaneous observation of a single general qualitative response and diverse quantitative responses to diets is coherent with the metabolic generalism hypothesis, and may help reconcile paradoxical notions about the consequences of generalism at the organismal level. One the one hand, individuals from generalist species do not always differ in response to various diets (e.g., *Leptinotarsa decemlineata*, Forister et al., 2007); or *Lymantria dispar*, Barbosa, 1978; Lazarevic et al., 1998). It is a defining characteristic of generalist species that they can tolerate (i.e., display relatively constant developmental responses under) a wider range of dietary circumstances when compared to specialist species. It therefore makes sense that the structural backbone of the metabolome of generalist individuals remains largely unaffected by diet, as uncovered here. On the other hand, the diverse quantitative responses of flies’ metabolomes to diet highlights the passive metabolic response of generalist *D. suzukii* individuals to the idiosyncrasies of their dietary environment. In particular, we observed that distinctive fruit secondary metabolites, relating to their specific flavor (flavonoids), color (anthocyanins) or scent (terpenoids), were accumulated in corresponding flies, and that very few *de novo* diet-specific fly metabolites were produced.

Our study thus suggests that diet generalization could stem from reduced care paid to hosts’ biochemical peculiarities and a neutral, passive stance towards potential hosts. While most of the ecological and evolutionary studies of host-plant interactions have focused on intricate plant-insect relations relying on strong selective forces, a passive and generic metabolic response could help species extend their diet breadth, at least transiently on evolutionary timescales. In this case, diet generalism would originate from a general tolerance rather than from the accumulation of independent genetic features associated to the exploitation of different resources (Forister et al., 2012; but see Calla et al., 2017), and the resulting plastic quantitative response to diet may not necessarily be adaptive.

Immediate costs of weakened defenses against specific plant compounds can be illustrated by the passive accumulation of terpenoids in blackcurrant-fed flies, indicating that *D. suzukii* may not have evolved efficient monoterpene metabolization pathways. In addition to their attractant or deterrent properties (Binder & Robbins, 1997), some terpenes have been shown to exhibit toxic properties and to be subjected to detoxification pathways in insects (Blomquist et al., 2021; Scalerandi et al., 2018). Indeed, in *D. suzukii*, a blackcurrant diet elicits a relatively large larval mortality and results in population extinction in only a couple generations (Olazcuaga et al., 2019, 2021). However, other fruits harboring potentially harmful compounds, such as amygdalin in cherry, can be excellent host for *D. suzukii*, including elevated oviposition preference and larval performance (Olazcuaga et al., 2019). Therefore, metabolic generalism in *D. suzukii* may not exclude some adaptations to cope with specific classes of toxic compounds, potentially reflecting its ancestral niche.

Metabolic generalism could also procure both immediate and longer-term benefits. For instance, accumulation of flavonoids increases the fitness of some phytophagous species (e.g., *Polyommatus icarus*; Simmonds, 2003). While chemical sequestration is an active process in specialist species, passive mechanisms may be common in generalist species (Petschenka & Agrawal, 2016). In the case of *D. suzukii*, the probably passive ‘sequestration’ of flavonoids might be useful against predators or parasitoids. On larger ecological scales, metabolic generalism could also entail benefits, e.g., by enabling a geographical expansion triggering enemy-release or by reducing intra-guild competition. In the case of *D. suzukii*, whose successive generations must exploit different hosts seasonally, an immediate reward of metabolic generalism could stem from resource allocation alternatives, e.g., a homeostatic immune response.

The passive accumulation of plant metabolites, such as anthocyanins, may also illustrate a greater opportunity of diet generalism due to plant domestication. Indeed, anthocyanins are known to trade-off with plant defense systems, and the domestication of fruit crops, by acting on anthocyanins accumulation, may have lowered plant defenses and thereby allowed generalist species to exploit niches that were ancestrally too challenging. For instance, it has been demonstrated that the domestication of cranberry cultivars, by selecting upon anthocyanin-related traits such as ripening and fruit color intensity, resulted in lowered plant defenses responses related to the jasmonic acid pathway (Rodriguez-Saona et al., 2011).

Finally, by showing that fruit-specific metabolites were readily found in fly metabolomes, our study suggests that metabolomic data might help track the diet of phytophagous insects in the wild (Alberdi et al., 2019; Nielsen et al., 2018). This could complement, high-throughput DNA sequencing-based diet reconstruction, that rely on the presence of foreign DNA material. Prerequisites for achieving this goal include an increased ability to correctly identify plant secondary metabolites and laboratory experiments to study the effect of various diets on the metabolomes of flies during development.

## Conclusion

This study is a first step toward a finer understanding of the mechanisms supporting the evolution of diet breadth using a joint metabolomic approach of diets and consumers, here underlining the importance of a neutral metabolic generalism. In the future, the investigation of different spatial and temporal scales will help to shed light on the equilibrium between diet generalism and specialization (Devictor et al., 2010), and on the origins and limits to diet generalism (Bolnick et al., 2003; Roughgarden, 1972; Sexton et al., 2017). To date, the interaction of phytophagous insect species such as leaf-chewing and mining insects with their host plants have drawn much of the research attention. Compared to more general plant tissues, fruits are specialized organs interacting with multiple species, with detrimental or beneficial impacts to plant fitness, and this may have shaped a greater complexity of chemical landscape in fruits. If the response to insect pests tends to decrease the attractiveness or palatability of fruits for more desirable seed dispersers, such as vertebrate frugivores, anti-insect defenses may perturb the reproductive potential of host plants (Andersen, 1988; but see Wilson et al., 2012). Understanding the ecological constraints exerted on both the chemistry of fruits and on the physiology of their consumers, impacting the evolution of niche breadth itself, will require a detailed knowledge of molecular- to community-level interactions across multiple trophic levels.

## Materials & Methods

### Fruit samples & media preparation

We investigated the influence of four major and economically important host plants of *D. suzukii*: cherry (*Prunus avium*, Rosaceae), cranberry (*Vaccinium macrocarpon*, Ericaceae), strawberry (*Fragaria x ananassa*, Rosaceae) and blackcurrant (*Ribes nigrum*, Grossulariaceae). To focus on their biochemical properties, we used artificial media based on industrial purees of these four fruits, instead of whole fruits, allowing to avoid effect of fruit skin, color, size and shape. Fruit purees were produced only from fruits and without added sugars, coloring or preservatives, by commercial companies (cherry: Huline S.A.S, Béziers, France; cranberry: Sicodis, Saint Laurent d’Agny, France; strawberry and blackberry: La Fruitière du Val Evel, Evellys, France). The use of industrial fruit purees enabled us to attain a homogenous fruit media composition regardless of ripening periods of each fruit.

To allow the development of *D. suzukii*, fruit media consisted of fruit puree, to which was added a simplified, protein-low version of the neutral medium classically used in drosophila laboratories (i.e., ‘German Food’ medium; https://bdsc.indiana.edu/information/recipes/germanfood.html). More precisely, the fruit media consisted of 60% of fruit puree and 40% of a mix of agar and yeasts, supplemented with ethanol and an antifungal compound (see Olazcuaga et al., 2022, for the full recipe).

### Flies

Our *Drosophila suzukii* stock population was initiated by sampling flies in the region of Montpellier (France) in September and October 2016. At this time of year and in this area, none of the four fruits studied in this study were available. Based on Poyet et al. 2015, flies could have potentially emerged from more than 18 wild host plants, which could be available in the sampling area in October (Table S1 in Olazcuaga et al. 2021). Before starting the experiment, this population was maintained at a size of ∼1000 individuals for a dozen generations in 10 ml flasks neutral medium (consisting of sugar, dry yeast, minerals, and antifungal solution; Backhaus et al., 1984), at 21 ± 2°C, 65% relative humidity, and a 16:8 (L:D) h light cycle in an air-conditioned chamber. Generations were non-overlapping and consisted of 24 hours of oviposition followed by 15 days of development and 6 days of sexual maturation and mating (for more details on the maintenance conditions of this population, see Olazcuaga et al. 2019).

### Diet experiment

To study the consequences of fruit diet on *D. suzukii*, we formed 4 populations of adults developed on a single fruit medium (cherry, cranberry, strawberry, and blackcurrant) from our stock population. To this aim, 20 groups of 20 six-day-old adults from the stock population (i.e., developed in the neutral medium) were left to mate and oviposit on a single fruit medium for 2 consecutive periods of 24h and then discarded. Ten days after oviposition and immediately after emergence, resulting adults were collected and maintained for seven days on the same fruit medium, with a maximum density of 40 adults per tube. For each fruit medium, these seven-day-old individuals were then flash frozen at - 80°C in groups of 100 individuals. For each fruit, three groups of 100 flies were stored at −80°C. This experiment was replicated in the exact same conditions for two consecutive generations of the stock population, in order to obtain biological replicates.

### Sample preparation

Frozen flies were freeze-dried for 24 hours. Each sample, composed of 30 flies, was weighed. Three samples were collected for each fruit diet and biological replicate (i.e., six samples for each diet), except for blackcurrant flies where only one biological replicate was available (i.e., three samples for blackcurrant flies). Flies were transferred into 2mL Eppendorf tubes and were crushed in a bead mill (TissueLyser II, Qiagen, Venlo, Netherlands) to obtain a powder. Metabolites were then extracted with methanol (20 μL/mg dry weight) containing 1 μg/mL of chloramphenicol as an internal standard. For fruit purees, 300 mg (fresh weight) were extracted with methanol (5 μL/mg fresh weight) containing 1 μg/mL of chloramphenicol. Extractions were performed in triplicate. Samples were placed in an ultrasonic bath (FB15050, Fisher Scientific, Hampton, USA) for 10 minutes, followed by centrifugation at 12000 g for 10 minutes at 4 °C. Supernatants (about 150 μL) were collected and transferred into vials for UHPLC-MS analyses.

### Non-targeted UHPLC-MS analyses

Extracts were analyzed using an UHPLC system (Dionex Ultimate 3000; Thermo Fisher Scientific) equipped with a diode array detector (DAD). Chromatographic separation was performed on a Nucleodur HTec column (150 × 2 mm, 1.8 μm particle size; Macherey-Nagel) maintained at 30 °C. The mobile phase consisted of acetonitrile/formic acid (0.1%, v/v) (eluant A) and water/formic acid (0.1%, v/v) (eluant B) at a flow rate of 0.25 ml/ min. The gradient elution program was as follows: 0–4 min, 80–70% B; 4–5 min, 70–50% B; 5–6.5 min, 50% B; 6.5–8.5 min 50–0% B; 8.5–10 min, 0% B. The injected volume of sample was 1μL. The liquid chromatography system was coupled to an Exactive Orbitrap mass spectrometer (Thermo Fisher Scientific) equipped with an electrospray ionization source operating in positive and negative modes. The instruments were controlled with Xcalibur software (Thermo Fischer). The ion transfer capillary temperature was set at 320 °C and the needle voltage at positive and negative modes at 3.4 kV and 3.0 kV, respectively. Nebulization with nitrogen sheath gas and auxiliary gas were maintained at 40 and 5 arbitrary units, respectively. The spectra were acquired within the mass-to-charge ratio (*m/z*) range of 95–1200 atomic mass units (a.m.u.), using a resolution of 50 000 at *m/z* 200 a.m.u. The system was calibrated internally using dibutyl phthalate as lock mass, giving a mass accuracy <1 ppm. LC-MS grade methanol and acetonitrile were purchased from Roth Sochiel (Lauterbourg, France), water was provided by a Millipore water purification system.

### Processing of metabolomic data

Raw data from all the samples of flies and fruit purees and from methanol as blank control, were converted to the mzXML format using MSConvert. mzXML data were sorted into nine classes as follows: four fruit-related data from four host plants (fruit purees), four fly-related data from groups of flies with host-specific diet, and blank. They were then processed using the XCMS software package (Smith et al., 2006). Ion detection parameters of the xcmsSet function of XCMS were as follows: “centWave”, ppm = 2, noise = 30 000, mzdiff = 0.001, prefilter = c(5,15000), snthresh = 6, peak width = c(6,35). Peaks were aligned using the obiwarp function using the following group density settings: bw = 10, mzwid = 0.0025, minimum fraction of samples for group validation: 0.5. Ion identifiers were generated by the XCMS script as MxxxTyyy, where xxx is the *m/z* ratio and yyy the retention time in seconds. For each ion identifier in each sample, an area corresponding to the integration of the MxxxTyyy peak was generated. All ions present in blank injections (areas in the blanks equal or sup. as those of samples) were suppressed from the datasets. This allowed the alignment of 7.805 peaks for the dataset in the positive mode and 3.665 peaks for the dataset in the negative mode. After specifying the detection mode for each ion (positive or negative), both datasets were merged. Finally, fruit-related and fly-related data were separated, resulting in a *fruit* dataset and a *fly* dataset.

### Statistical analyses

All statistical analyses were performed using R v3.6.1 (R Core Team, 2019).

### Non-targeted metabolomic analysis of fruits

To ensure that the selected fruit purees did have different compositions, we performed a Principal Component Analysis (PCA) on the scaled ion quantities of the *fruit* dataset using the *FactoMineR* R package (Lê et al., 2008). We additionally plotted the heatmap of the two-dimensional clustering analysis of centered and scaled metabolite quantities of the *fruit* dataset using the *ComplexHeatmap* R package (Gu et al., 2016). Seriations of ions (rows) and fruit samples (columns) were performed using the “traveling salesperson” and the optimal leaf ordering methods of the *seriation* R package, respectively (Hahsler et al., 2020).

### Host use

As a preliminary step, we checked whether ions content of flies was due to direct contamination by fruit by investigating the global correlation between the quantities of ions in fruit purees and in flies that developed on them. Presence of a correlation would be indicative of contamination, while the absence of correlation would reveal an absence of contamination of flies by the purees.

To check whether *D. suzukii* flies consume different fruits in a similar (i.e., ‘metabolic generalism’) or in a dissimilar way (i.e., ‘multi-host metabolic specialism’), we used both the *fruit* and *fly* (quantitative) datasets and a presence/absence (qualitative) dataset derived from them. The presence/absence dataset was constructed as follows: all metabolites that were present in at least three replicates were considered present, otherwise they were considered absent. Three types of analyses used this presence/absence dataset.

First, we computed intersection sizes between all fruits’ and flies’ ions from the presence/absence dataset considered together and investigated these intersections using upset plots (bar plots designed to represent intersections; *UpSetR* R package, Conway et al., 2017). Indeed, in the case of a similar use of different diets (metabolic generalism), flies should all share a large part of their ions, fly ions specific of a particular diet should be rare or absent, and/or ions specific of single fruit-fly pair (i.e., a pair of fruit and flies grown over the same fruit) should also be rare (Figure 1C). Reciprocal results would favor the multi-host metabolic specialism hypothesis.

Second, we separately computed the intersections of ions present in all fruits on one hand and in all flies on the other hand. We then related these two datasets to examine (*i*) whether ions most common to all fruits are also common to all flies, (*ii*) whether ions unique to a given fruit are also unique to flies that developed on the same fruit, and (*iii*) what is the origin of diet-specific fly ions. Results of these computations were plotted using the *ggalluvial* R package (Cory & Read, 2020). In this analysis, differently-fed flies metabolizing ions that are shared between fruits to produce ions shared between flies would be indicative of metabolic generalism, while differently-fed flies metabolizing ions private to a given fruit to produce new ions (mostly not shared between flies) would be indicative of multi-host metabolic specialism (Figure 1E).

Finally, we visually investigated the multi-dimensional clustering of fruits’ and flies’ metabolomes using a global PCA with the (quantitative) *fruit* and *fly* datasets (Figure 1F). Metabolic generalism ought to cause all flies’ metabolomes to cluster together, regardless of diet, while multi-host metabolic specialism would be detected if fruit-fly pairs of metabolomes cluster together (Figure 1G).

### Metabolomic discrimination of flies according to their diets

To investigate the putative discrimination of flies according to their diet, we first conducted a PCA analysis on the scaled ion quantities of the large *fly* dataset, using the *FactoMineR* R package (Lê et al., 2008). Following the results of the previous PCA (see Results section), we computed a list of fly ions most associated with each diet using generalized linear models with Elastic Net regularization (Z ou & Hastie, 2017), as implemented in the *glmnet* R package (Friedman et al., 2010). Elastic Net regularization is particularly useful to distinguish between causal features (in our case, ions) between treatments when the number of samples is small in comparison with the number of features (Kirpich et al., 2018). The Elastic Net regularization combines (in proportion *α*) LASSO and ridge regression methods to build on their respective strength and detect significant features in highly correlated spaces, such as found in ∼omics data (Acharjee et al., 2013; Gonzales & Saeger, 2018; Waldmann et al., 2013). For each diet, all fly samples were used and classified as either from the given diet or not. Preliminary exploration of Elastic Net mixing parameter *α* and regularization parameter λ using 10-fold cross validation with the *caret* R package (Kuhn, 2021) indicated that the method provided good sample classification using various combinations of *α* and λ. Two sets of parameters were then used following two distinct goals. The ‘large’ set of parameters (*α* = 0.5 and λ = 0.01, except λ_blackcurrant_ = 0.001) was constructed to enable optimal classification of flies according to diets. The ‘compact’ set of parameters (*α* = 0.9 and λ = 0.001, except λ_blackcurrant_ = 0.0001) was modified to enable manual curation of the list of metabolites significantly associated to diet, by reducing its overall length.

We plotted the heatmap of the two-dimensional clustering analysis of scaled quantities of the flies’ ions present on the ‘large’ list for both flies and fruits to simultaneously visualize (*i*) the classification performance of flies according to their diet and (*ii*) the relative quantities of these flies’ ions in every fruit, using the *ComplexHeatmap* R package (Gu et al., 2016). We tested whether ions of the ‘large’ list for a given diet were significantly more present in the flies of the same diet than in flies of alternate diets, and in the same fruit than in alternate fruits. To this aim, we used a bootstrap resampling procedure (n = 1,000 replicates) to estimate median quantities of ions in all flies and fruits and their 95% confidence intervals, as recommended for heavily skewed continuous data containing zero values (Helsel, 2011). For each diet, ion quantities in flies were plotted against the ion quantities in their respective fruit (on a log scale), and their correlation (and associated R²) computed through linear modelling. Additionally, we investigated the intersections of origin (and their degrees) of these diet-specific ions, both in fruits and flies, to gain insight about their specificity.

Finally, to evaluate the respective power of our qualitative and quantitative datasets to correctly classify flies according to their diet, we randomly sampled ions in batches of various sizes (10 to 10,000 ions; each random sampling repeated 10,000 times). We then computed (*i*) the proportion of flies correctly classified to their diet and (*ii*) Adjusted Mutual Information (AMI, a comparison index between clustering; Vinh et al., 2010) for each sample size for both quantitative and qualitative datasets. AMI was computed using the *aricode* R package (Chiquet et al., 2020).

### Identification of metabolites

We attempted to identify two different categories of metabolites. First, we tried to identify metabolites from the compact’ list of ions associated to diet in flies (from our Elastic Net analysis). Second, we investigated 50 major ions in each fruit (*i.e.,* with the highest peak area), potentially corresponding to major metabolites in each fruit.

Metabolites were identified based on molecular formula generated from accurate mass measurements, given the fact that mass spectrometry conditions were chosen to obtain fragments by in-source fragmentation, in addition to pseudo-molecular ions. Putative metabolite identifications were proposed based on expertized analysis of the corresponding mass spectra and fragmentation patterns, in comparison with published literature. This has allowed the putative identification of 71 food-associated metabolites detected in flies, 26 identifications being confirmed with authentic standards. Finally, for each identified compound in each fruit, a survey of the published literature was performed to check for previously identification of this compound in this fruit. Standards for metabolite identification were purchased from Sigma-Aldrich (Saint-Quentin Fallavier, France) and Extrasynthese (Lyon, France).

Two-dimensional clustering analyses of the scaled quantities of identified compounds were computed overall fruit and fly samples, using ions selected from the ‘compact’ list and major fruit ions, and visualized using the *ComplexHeatmap* R package (Gu et al., 2016).

## Acknowledgments

JF was supported by recurrent institutional funding from INRAE. JF would like to thank Aude Gilabert, Pierre E. Sirius and Ariane N. Vega for their outstanding support. L.O. acknowledges support from the European Union program FEFER FSE IEJ 2014-2020 (project CPADROL), the INRAE scientific department SPE (AAP-SPE 2016) and the US National Science Foundation (DEB-1930650 to Ruth Hufbauer and Brett Melbourne).

## Figure Supplements

**Figure 1—figure supplement 1:**
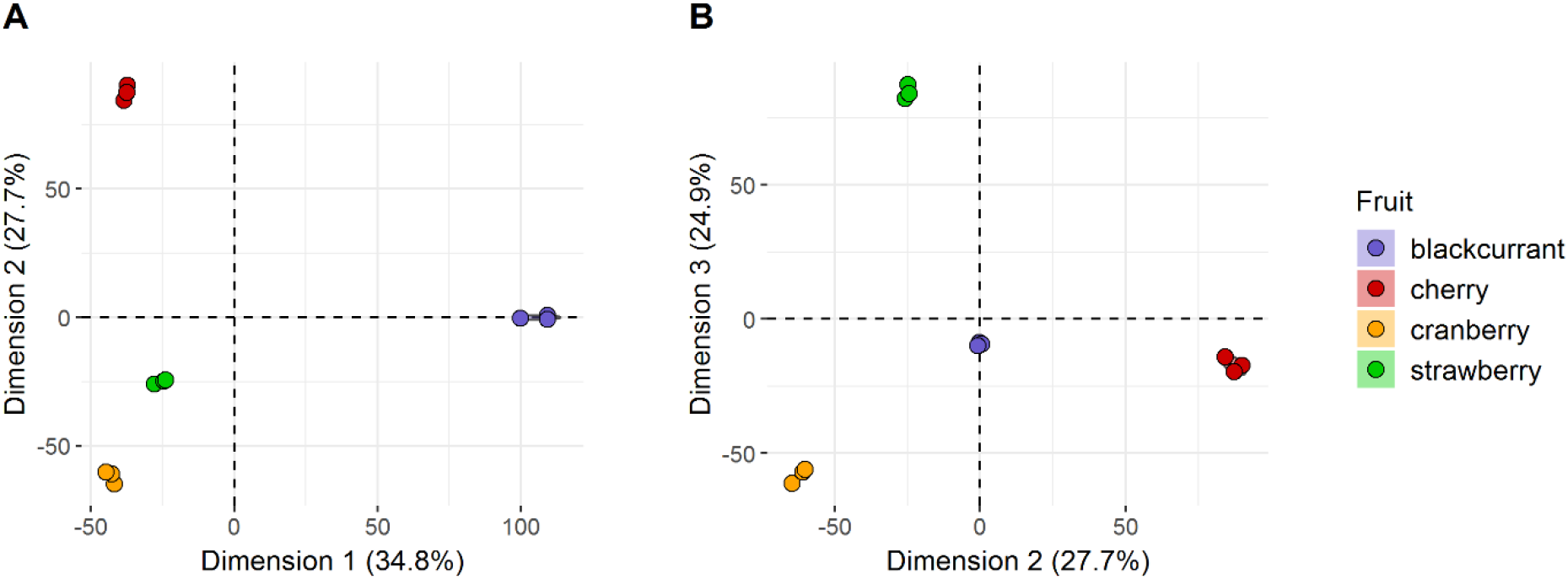
Principal component analysis (PCA) of fruits’ metabolomes. (A) Dimension 1 vs dimension 2, and (B) dimension 2 vs dimension 3. Different fruit samples are represented by colored dots. Eigenvalues of the PCA of fruit metabolomes showed that only three dimensions explained more than 87% of the dataset variance, thus we visually explored only the first three dimensions of the PCA.

**Figure 1—figure supplement 2:**
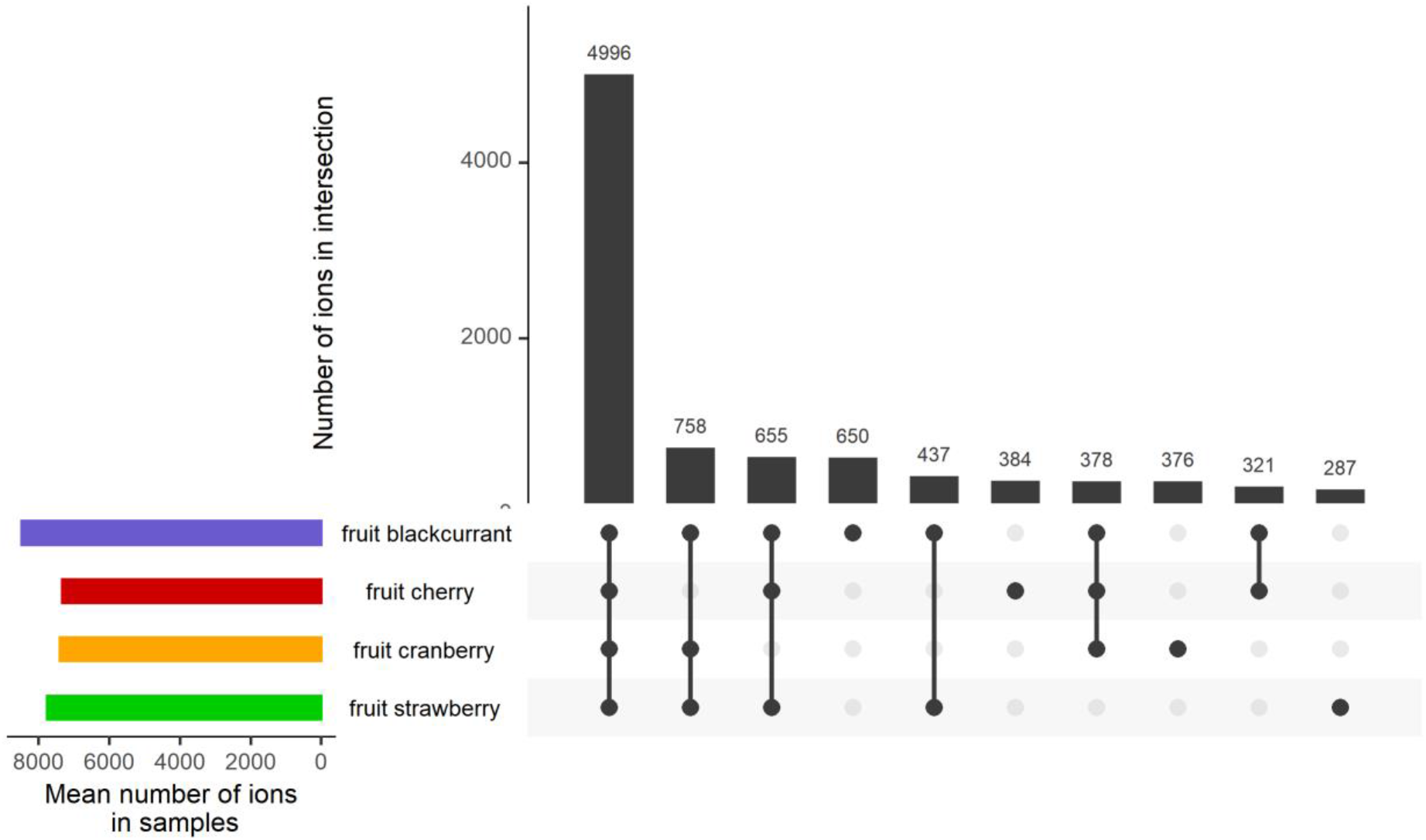
Upset plot of the number of shared ions between all fruits (ten largest intersections).

**Figure 1—figure supplement 3:**
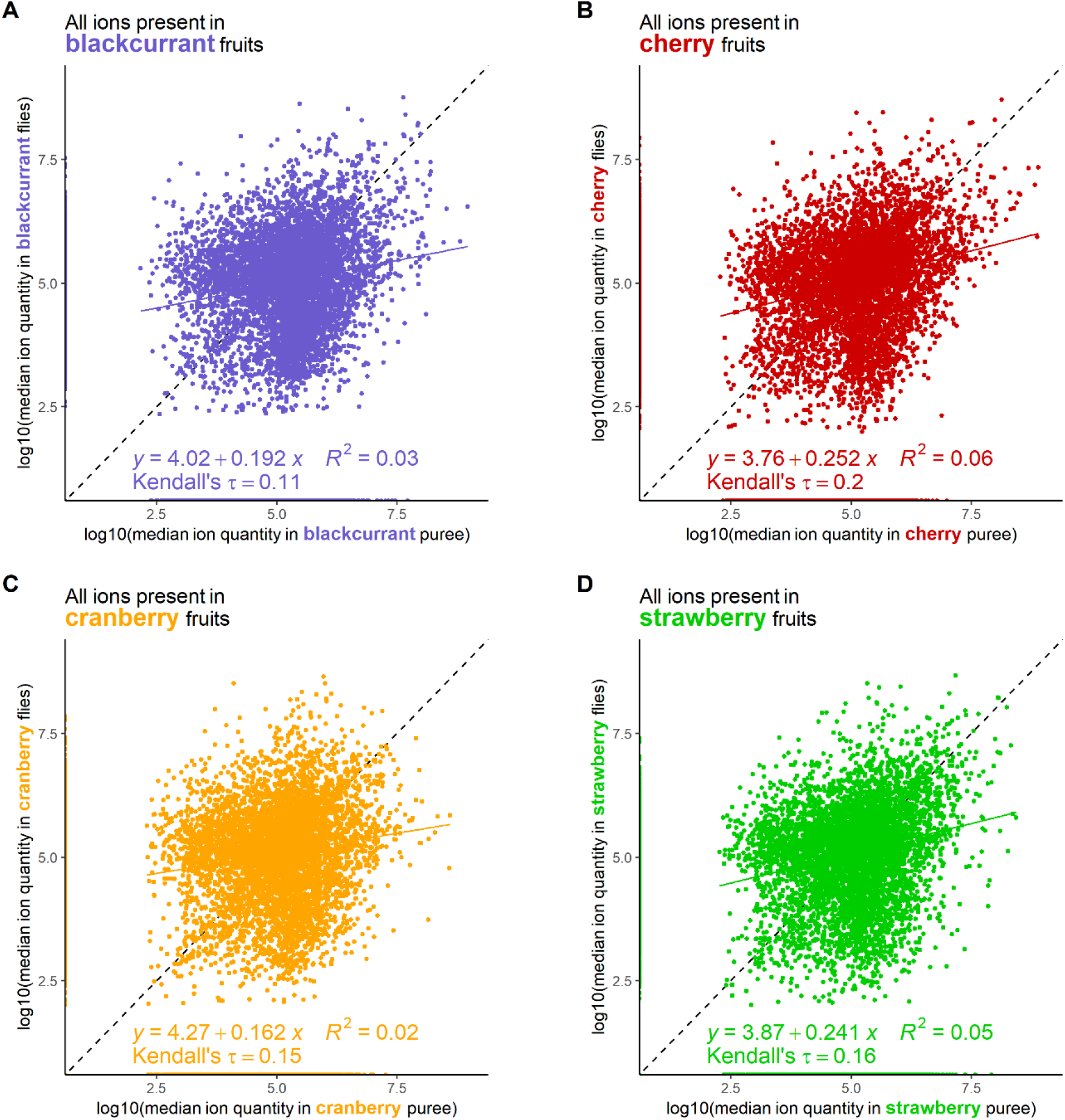
Relation between ion quantity in fruit vs in flies that developed on the same fruit. (A) blackcurrant, (B) cherry, (C) cranberry, (D) strawberry. Linear regression coefficients, R-square and Kendall’s τ are indicated for each fruit-fly pair.

**Figure 3—figure supplement 1:**
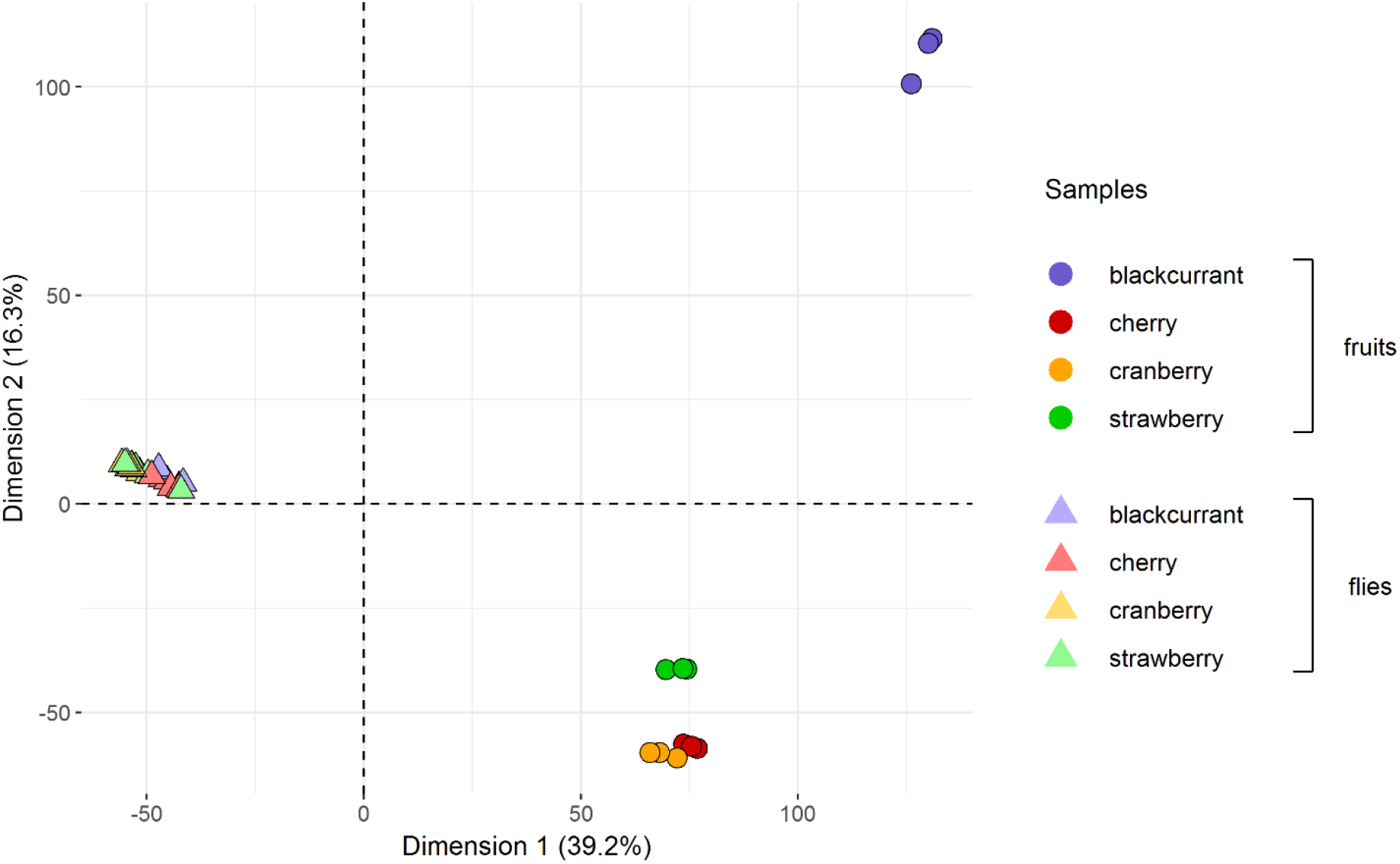
Fruits and flies’ metabolomes cluster separately and do not form pairs. Fruit and fly samples in this global Principal Component Analysis (PCA) are represented by dots and triangles, respectively. Colors represent different fruits or diets in the case of flies. All flies’ samples cluster together and away from fruits, regardless of their diets. See Figure 1G for expectations according to diet generalization hypotheses.

**Figure 4—figure supplement 1:**
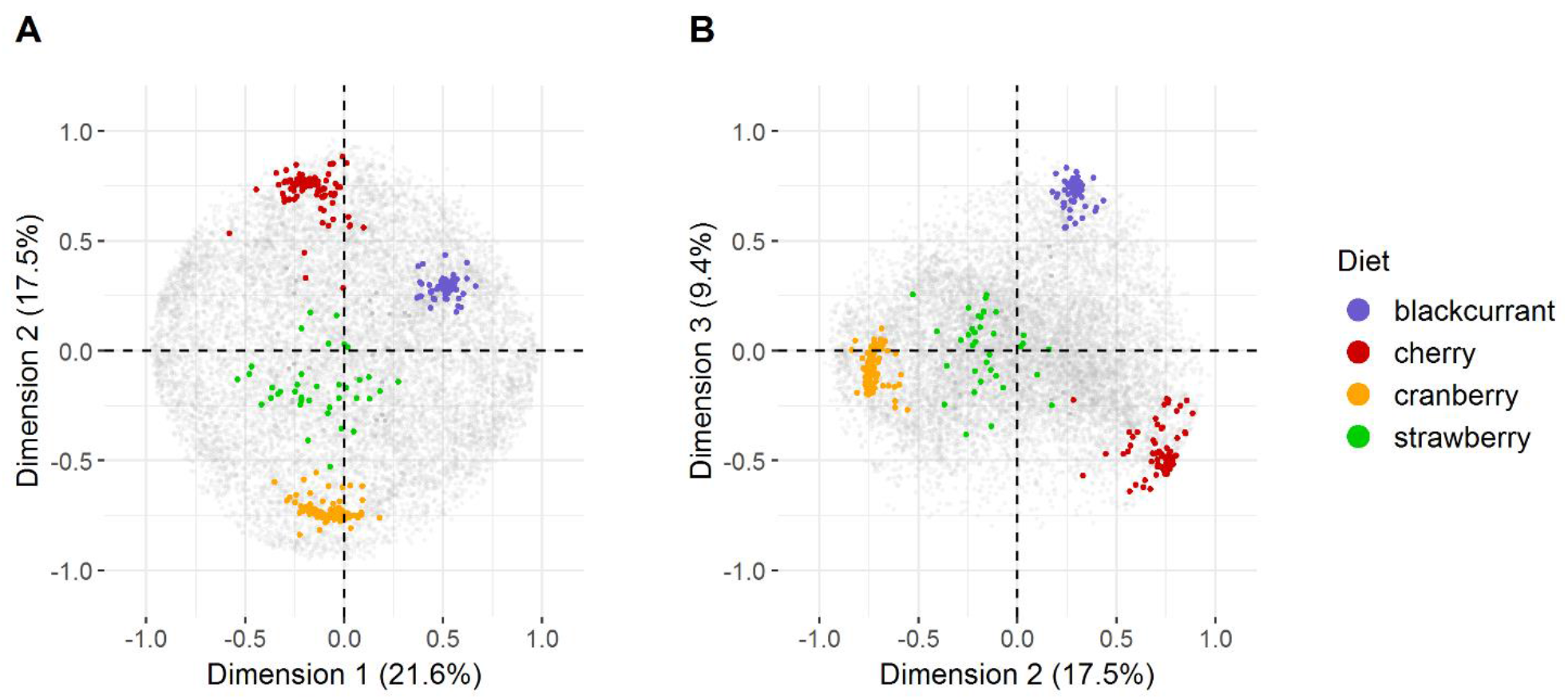
Plots of all individual ions of the Principal Component Analysis of fruit metabolomes. (A) Dimension 1 vs dimension 2, and (B) dimension 2 vs dimension 3. Fruit-specific ions are indicated in color, all other ions are indicated in gray.

**Figure 4—figure supplement 2:**
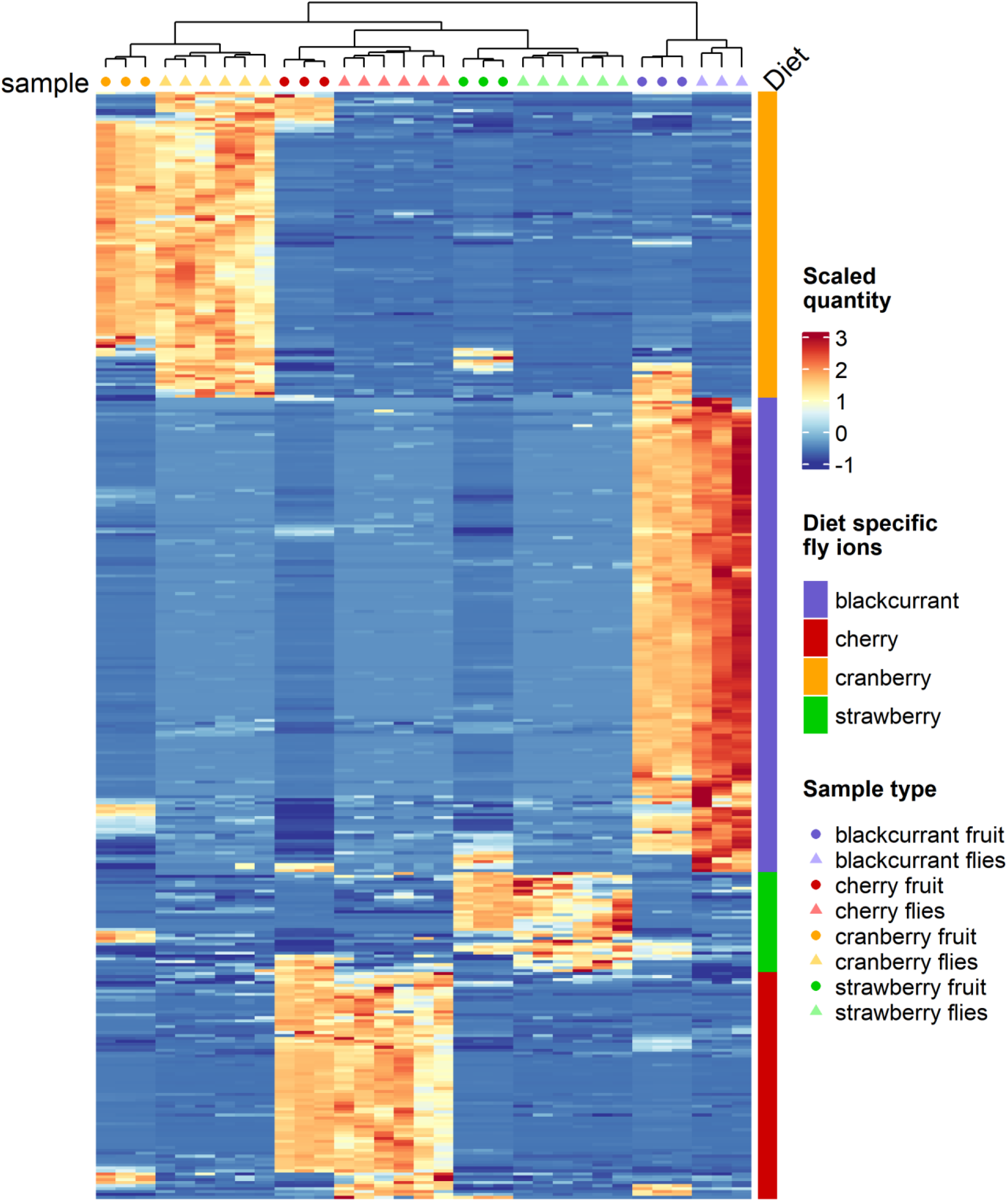
Classification of fruit and fly samples following selection of diet-specific ions through GLM with Elastic Net regularization. Heatmap scale colors indicate the relative quantity of each ion (row) of the ‘large’ list of diet-specific ions (specificity indicated by discrete colors; see main text for details).

**Figure 4—figure supplement 3:**
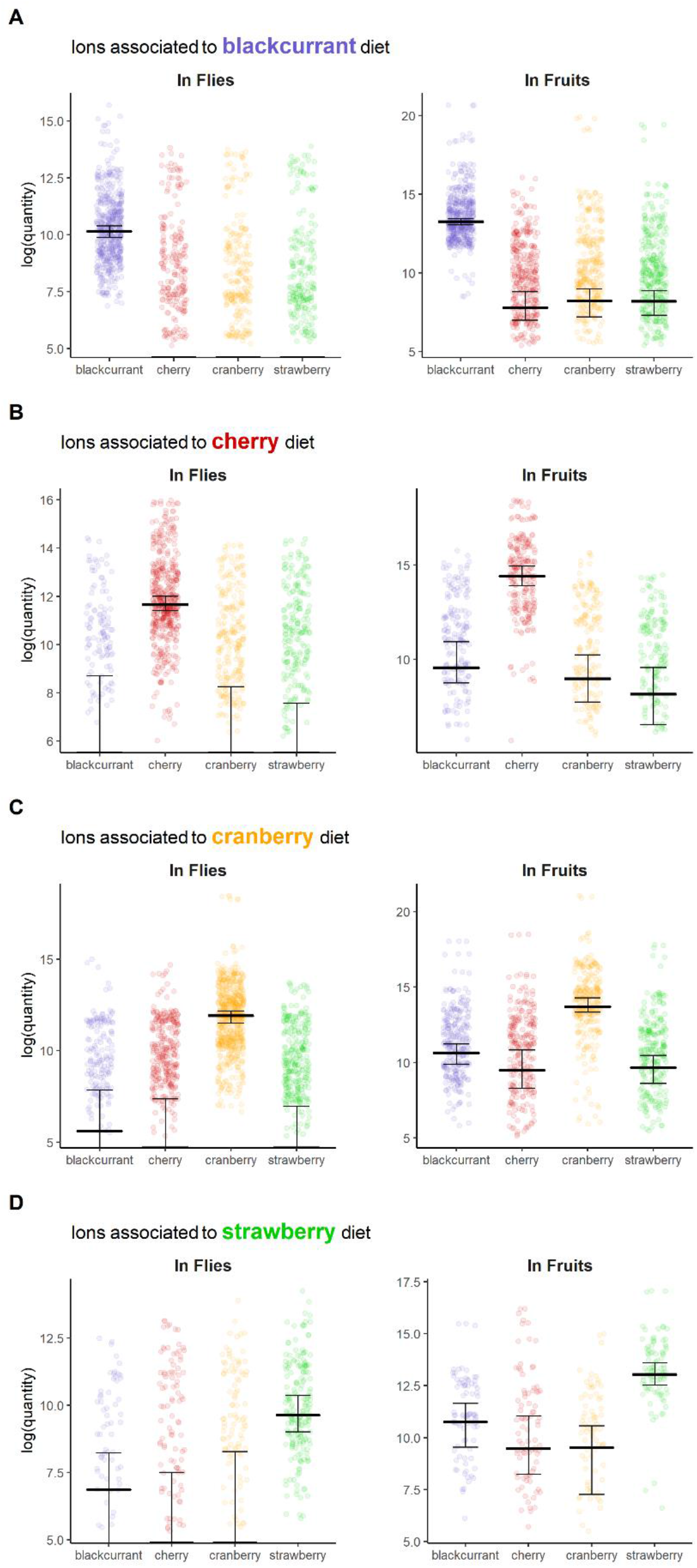
Quantitative levels of diet-specific fly ions in flies and fruits. (A) blackcurrant, (B) cherry, (C) cranberry, (D) strawberry. Medians and their confidence intervals are represented by black bars.

**Figure 4—figure supplement 4:**
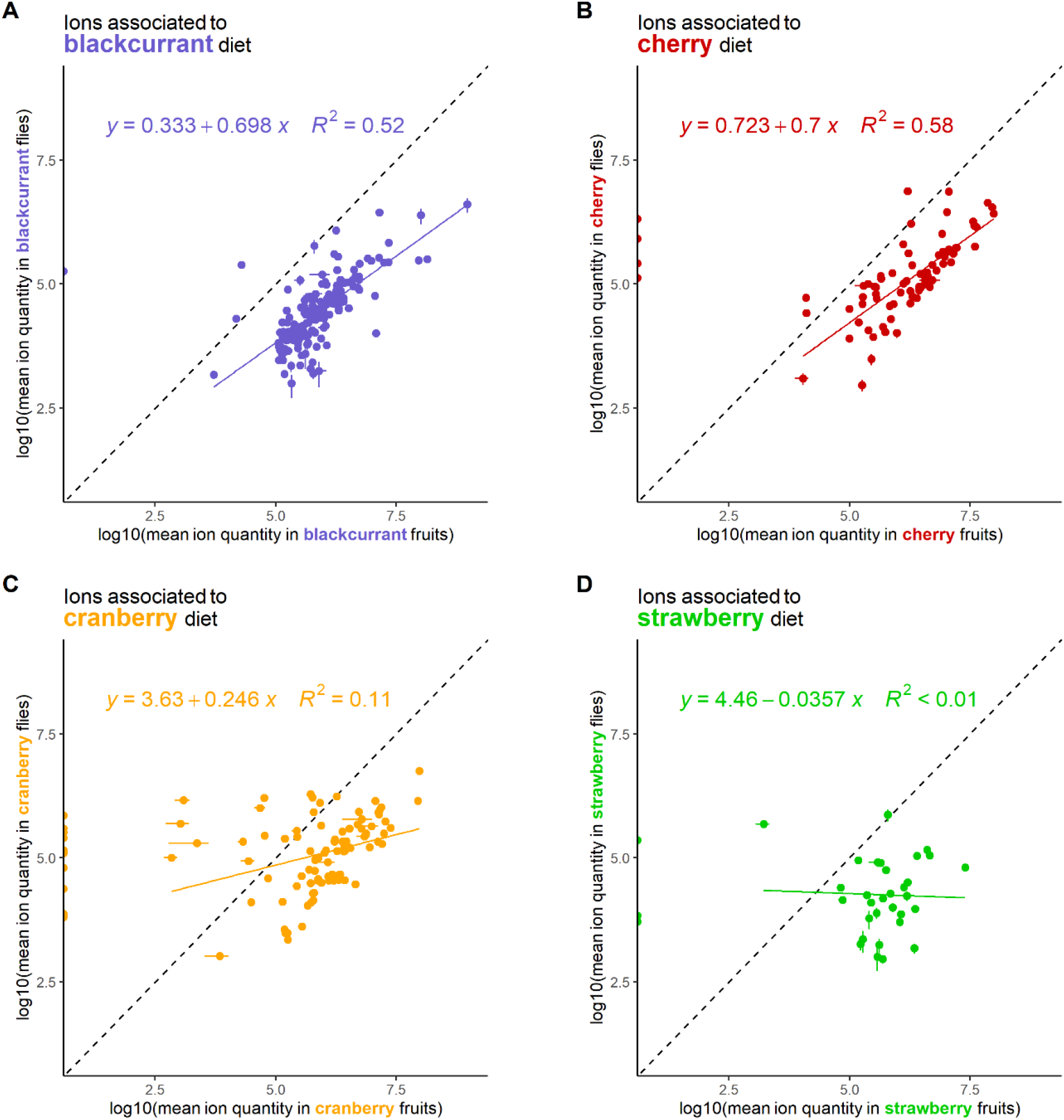
Relationships between the quantities of diet-specific fly ions in fruit and flies of the following fruit-fly pairs. (A) blackcurrant, (B) cherry, (C) cranberry, (D) strawberry.

**Figure 4—figure supplement 5:**
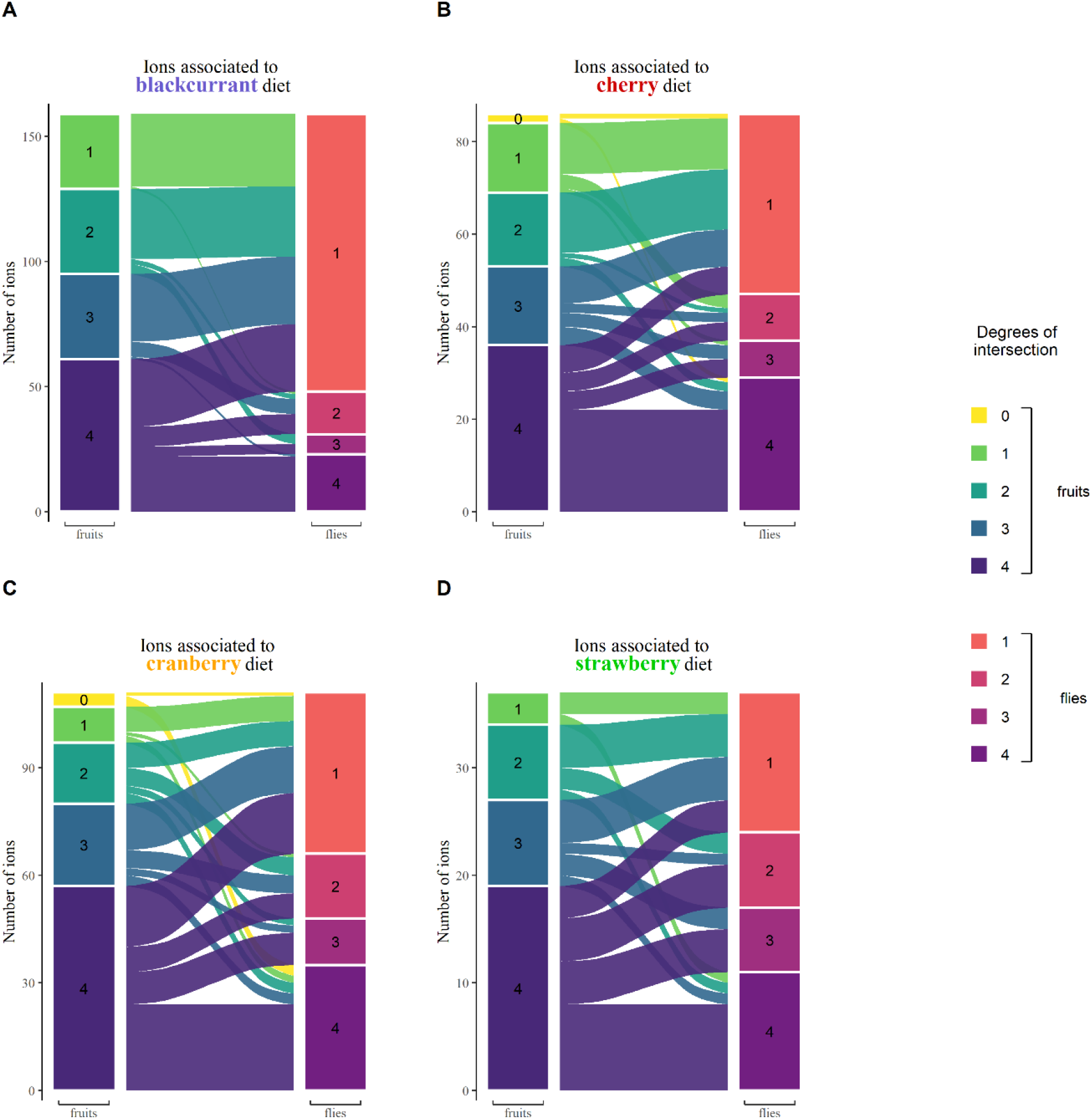
Intersection sizes (in fruits and flies) and relationships of fly ions specific of the following diets. (A) blackcurrant, (B) cherry, (C) cranberry, (D) strawberry. Ions that are shared within fruits or within flies show a higher degree of intersection.

**Figure 4—figure supplement 6:**
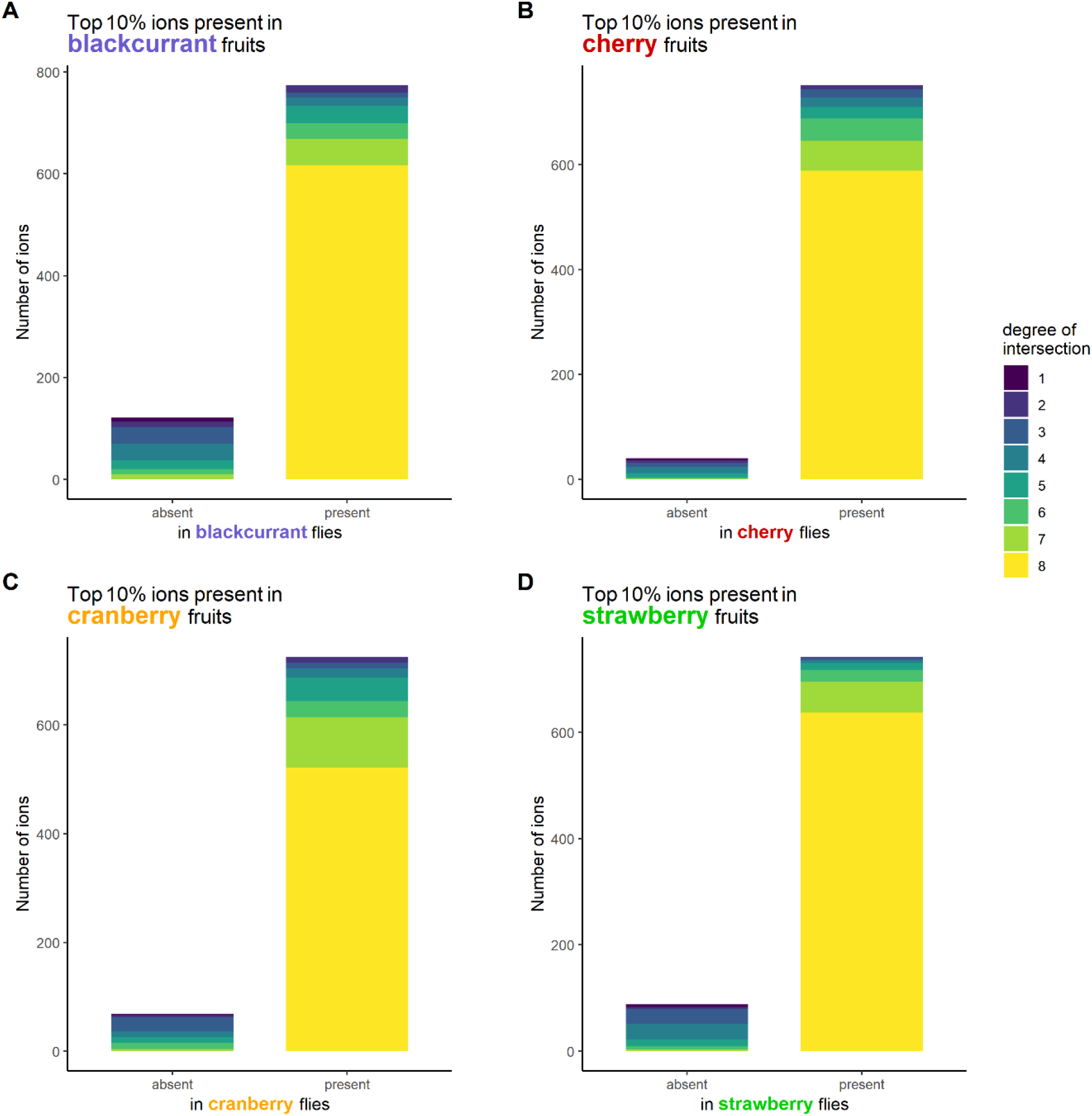
Number and degree of intersection of major fruit ions relative to their presence in the consumer flies. (A) blackcurrant, (B) cherry, (C) cranberry, (D) strawberry.

**Figure 4—figure supplement 7:**
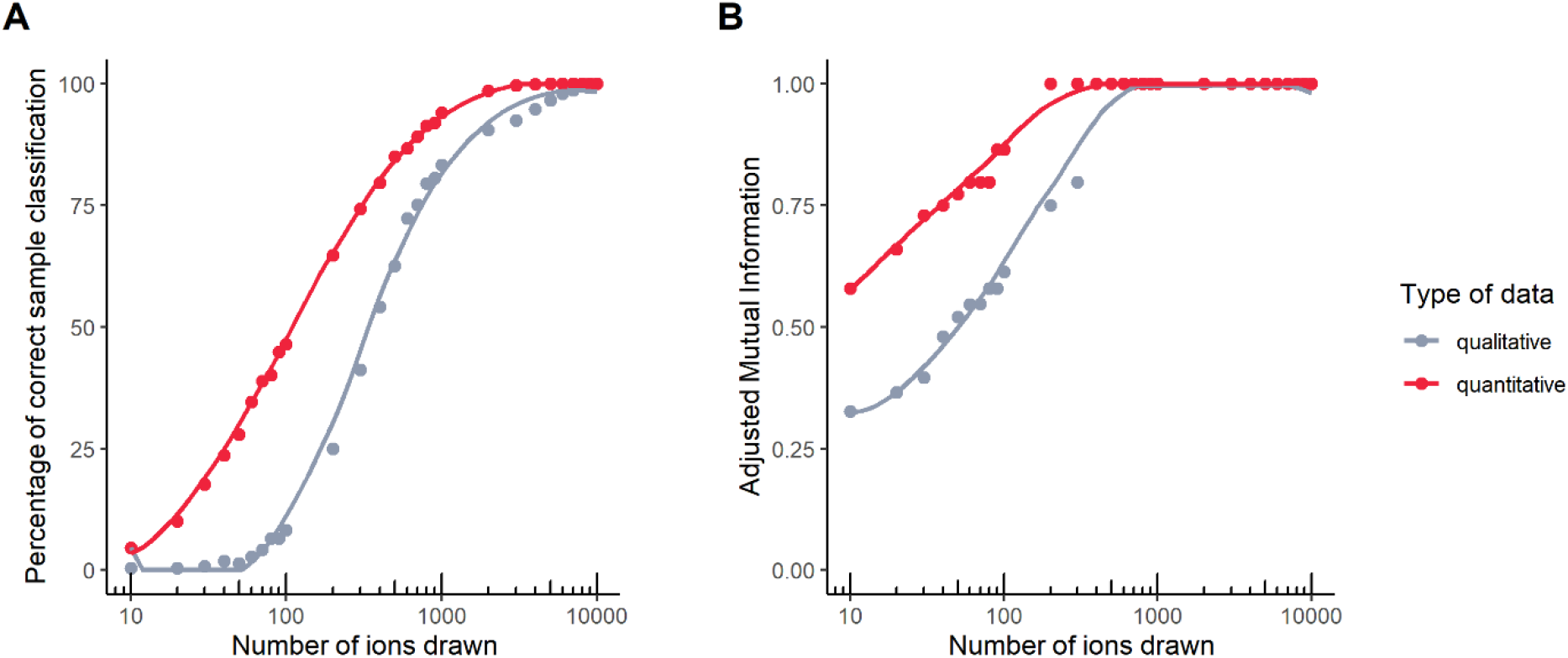
Classification performance of qualitative vs quantitative fly datasets to infer diet.: (A) percentage of correct classification, and (B) Adjusted Mutual Information.

**Figure 5—figure supplement 1:**
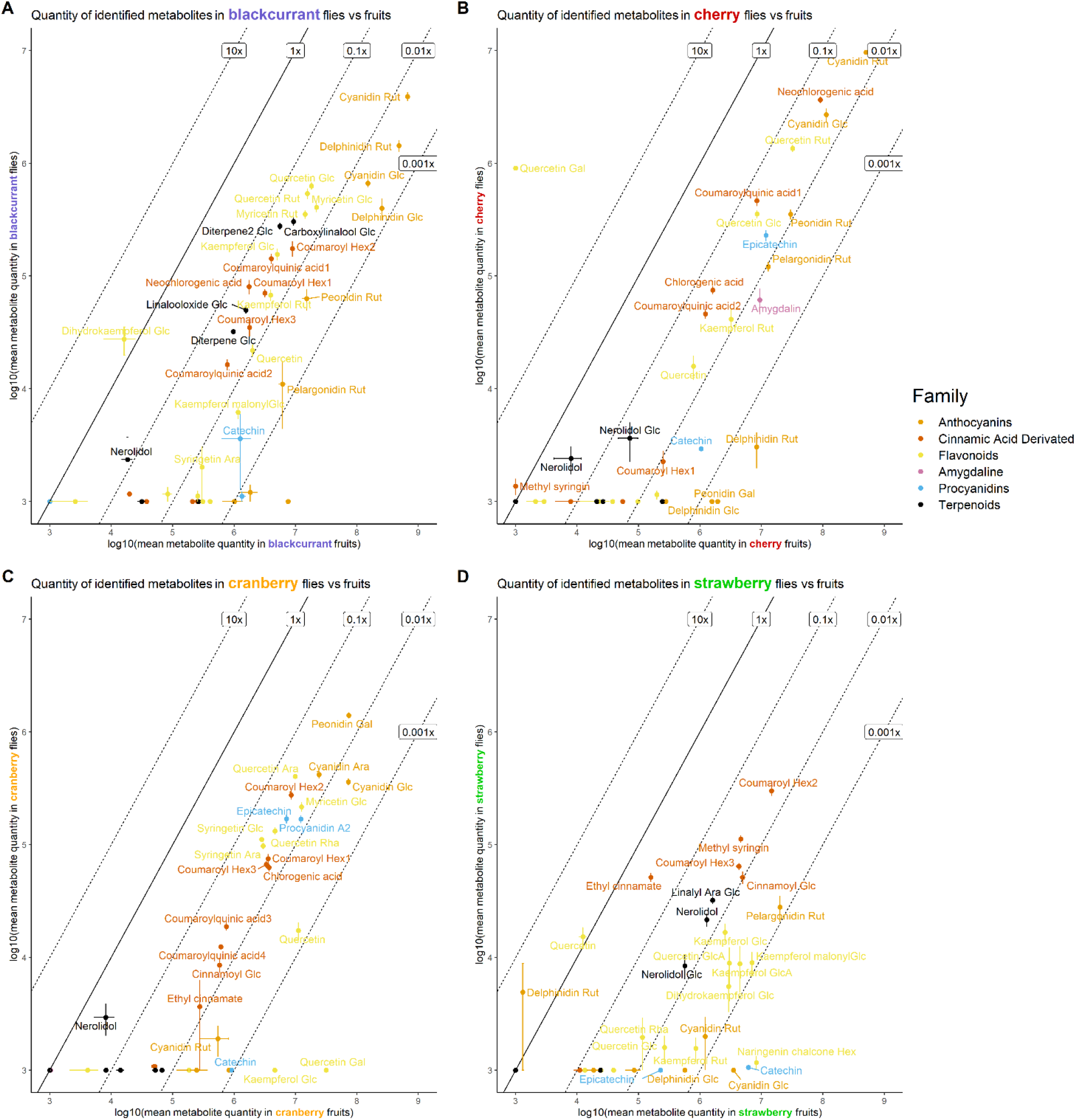
Quantities of identified metabolites in the following fruit-fly pairs. (A) blackcurrant, (B) cherry, (C) cranberry, (D) strawberry. Most metabolites are present ten to a hundred times less in flies that in their respective diet.

