## Supplementary Table 1 for "Metabolic consequences of various fruit-based diets in a generalist insect species"

List of the 71 identified metabolites

Abbreviations for chemical family names are as follows: AA, amino-acid; AN, anthocyanin; F, flavonoid; CAD, cinnamic acid derivated; BAD, benzoic acid derivated; OA, organic acid; PC, procyanidin; S, sugar; T, terpenoids; M, other metabolite. Abbreviated column names are as follows: M, [M+H]+ ion mass; RT, retention time; val, validation with commercial standard. The “flies” column indicates diet-specific fly metabolites present on the ‘compact’ list of fly ions (see Materials & Methods), and for which diet they have been detected as specific. The “fruits” column indicates metabolites the presence in one or several lists of top 50 fruit ions (in term of average quantity). Columns named after specific fruits indicate the references linking metabolites to the said fruit chemical composition. Full data of the mentioned literature can be downloaded as a XXX file to the following URL: XXXXXXXXXX

| **name** | **family** | **formula** | **M** | **Ionisation mode** | **RT** | **val** | **flies** | **fruits** | **blackcurrant** | **cherry** | **cranberry** | **strawberry** |
| --- | --- | --- | --- | --- | --- | --- | --- | --- | --- | --- | --- | --- |
| Asparagine | AA | C4H8N2O3 | 133,0608 | pos | 1,15 | yes |  | cherry, strawberry |  |  |  |  |
| Glutamine | AA | C5H10N2O3 | 147,0763 | pos | 1,15 | yes |  | blackcurrant, strawberry |  |  |  |  |
| Oxoproline | AA | C5H8N2O3 | 130,0499 | pos | 1,15 | no |  | strawberry |  |  |  |  |
| Proline | AA | C5H9NO2 | 116,0707 | pos | 1,18 | yes |  | cherry |  |  |  |  |
| Pelargonidin rutinoside | AN | C27H31O14 | 579,1707 | pos | 1,68 | no |  | strawberry | [1–3], but see [4] | [5–10] |  | [11–17] |
| Cyanidin glucoside | AN | C21H21O11 | 449,1078 | pos | 1,63 | yes | blackcurrant | blackcurrant, cherry, cranberry | [4,17–24] | [5–8,10,25–28] | [29–32] | [14–17,33] |
| Cyanidin arabinoside | AN | C20H19O10 | 419,0972 | pos | 1,68 | no | cranberry | cranberry | [2] |  | [29–32] |  |
| Cyanidin rutinoside | AN | C27H31O15 | 595,1658 | pos | 1,62 | no |  | blackcurrant, cherry | [4,17–24,34] | [5–8,10,25–28] |  | [15,16] |
| Cyanidin coumaroyl glucoside | AN | C30H27O13 | 595,1442 | pos | 3,61 | no |  |  | [2] |  |  |  |
| Peonidin galactoside | AN | C22H23O11 | 463,1234 | pos | 1,69 | no | cranberry | cranberry |  |  | [29,30,32,35] |  |
| Peonidin rutinoside | AN | C28H33O15 | 609,1813 | pos | 1,68 | no | cherry | cherry | [2,22] | [5–8,10,25–28] |  |  |
| Delphinidin glucoside | AN | C21H21O12 | 465,1026 | pos | 1,26 | yes | blackcurrant | blackcurrant | [1,4,17,18,20–24,34] |  |  |  |
| Delphinidin rutinoside | AN | C27H31O16 | 611,1603 | pos | 1,32 | no |  | blackcurrant | [1,4,17–24,34] |  |  |  |
| Naringenin chalcone hexoside | F | C21H20O10 | 433,1129 | pos | 5,62 | no |  |  |  |  |  | [36] |
| Kaempferol glucoside | F | C21H30O11 | 459,1861 | pos | 5,05 | yes | blackcurrant |  | [1,20,22,37,38] | [8,26] |  | [16,33,39] |
| Kaempferol glucuronide | F | C21H18O12 | 463,0871 | pos | 5,32 | yes |  |  |  |  |  | [33] |
| Kaempferol rutinoside | F | C27H30O15 | 595,1659 | pos | 4,48 | yes |  |  | [22,38] | [26,27,40] |  |  |
| Kaempferol malonylglucoside | F | C24H22O14 | 535,1082 | pos | 5,95 | no |  |  |  |  |  | [16,39] |
| Dihydrokaempferol glucoside | F | C21H22O11 | 451,1235 | pos | 2,67 | no |  |  |  |  |  | [16,33,39] |
| Quercetin | F | C15H10O7 | 303,0498 | pos | 7,16 | yes |  |  | [4,41,42] |  | [43–47] | [33] |
| Quercetin rhamnoside | F | C21H20O11 | 447,0930 | neg | 6,28 | yes |  | cranberry |  | [8] | [47] |  |
| Quercetin arabinoside | F | C20 H18 O11 | 435,0922 | pos | 4,92 | no | cranberry | cranberry |  |  | [45,46] |  |
| Quercetin glucoside | F | C21H20O12 | 465,1027 | pos | 4,14 | yes |  | blackcurrant, cherry | [20,22,23,38] | [8,26,40] | [47] | [16,36,39] |
| Quercetin glucuronide | F | C21H18O13 | 479,0819 | pos | 4,52 | yes |  |  | [38] |  |  | [16,39] |
| Quercetin rutinoside | F | C27H30O16 | 611,1608 | pos | 3,6 | yes |  | blackcurrant, cherry | [20,22,23,37] | [8,25–28,40] |  | [16,39] |
| Quercetin galactoside | F | C21H20O12 | 463,0881 | neg | 5,22 | no | cherry | cranberry | [38] | [8] | [45–47] |  |
| Myricetin glucoside | F | C27H30O17 | 481,0977 | pos | 3,15 | no |  | blackcurrant | [22,23] |  |  |  |
| Myricetin rutinoside | F | C27H30O17 | 627,1555 | pos | 2,8 | no | blackcurrant | blackcurrant | [22,23,37] |  |  |  |
| Syringetin arabinoside | F | C22H22O12 | 479,1184 | pos | 6,37 | no |  |  |  |  |  |  |
| Syringetin glucoside | F | C23 H24 O13 | 507,1138 | neg | 4,98 | yes | cranberry |  |  |  |  |  |
| Coumaroylquinicacid1 | CAD | C16H18O8 | 337,0926 | neg | 2,33 | no | cherry | cherry | [22] | [5–8,10,25,27] |  |  |
| Coumaroylquinic acid2 | CAD | C16H18O8 | 337,0927 | neg | 3,32 | no |  |  | [22] | [5–8,10,25,27] |  |  |
| Coumaroylquinic acid3 | CAD | C16H18O8 | 337,0927 | neg | 3,03 | no |  |  | [22] | [5–8,10,25,27] |  |  |
| Coumaroylquinic acid4 | CAD | C16H18O8 | 337,0927 | neg | 3,68 | no |  |  | [22] | [5–8,10,25,27] |  |  |
| Neochlorogenic acid | CAD | C16H18O9 | 353,0876 | neg | 1,88 | yes | cherry | cherry | [20,22] | [5–8,10,25] |  |  |
| Chlorogenic acid | CAD | C16H18O9 | 355,1076 | pos | 2,8 | yes |  | cranberry | [20,22] | [5–8,10,25] | [48,49] |  |
| Cinnamoyl glucoside | CAD | C16H20O9 | 355,1033 | pos | 5,2 | no |  | strawberry |  |  |  |  |
| Coumaroyl Hexoside1 | CAD | C15H18O8 | 325,0926 | neg | 2,18 | no |  |  | [50] |  | [50,51] | [50] |
| Coumaroyl Hexoside2 | CAD | C15H18O8 | 325,0925 | neg | 2,3 | no | strawberry |  | [50] |  | [50] | [50] |
| Coumaroyl Hexoside3 | CAD | C15H18O8 | 325,0925 | neg | 2,5 | no |  | strawberry | [50] |  | [50] | [50] |
| Methyl syringin | CAD | C18H26O9 | 385,1502 | neg | 7,13 | no | strawberry |  |  |  |  | [52] |
| Amygdalin | M | C20H27O11N | 475,1922 | pos | 2,27 | no |  | cherry |  |  |  |  |
| Indole derivated | M | C13H14N2O2 | 231,1128 | pos | 2,38 | no | blackcurrant |  |  |  |  |  |
| 3-carboxymethyl-Indole-N-glucoside | M | C16H19O7N | 336,1086 | neg | 2,92 | no | blackcurrant |  | [53] |  |  |  |
| Ethyl cinnamate | CAD | C11H12O2 | 177,0909 | pos | 7,64 | yes | strawberry |  |  |  |  | [54] |
| Acylphloroglucinol glucoside | M | C16H22O9 | 359,1336 | pos | 5,3 | no | strawberry |  |  |  |  | [55] |
| Benzylalcohol diglucoside | BAD | C19H28O11 | 431,1560 | neg | 1,95 | no | cherry |  |  | [56] |  |  |
| Glucuronic Acid | OA | C6H10O7 | 193,0345 | neg | 1,28 | yes |  | blackcurrant |  |  |  |  |
| Gluconic acid | OA | C6H12O7 | 195,0504 | neg | 1,3 | yes |  | cherry, strawberry |  |  |  |  |
| Apionic acid | OA | C5H10O6 | 165,0397 | neg | 1,27 | no |  | blackcurrant |  |  |  |  |
| Ascorbic acid | OA | C6H8O6 | 175,0240 | neg | 1,32 | yes |  | blackcurrant | [19,38,42] | [57] | [29,30] | [14,58] |
| Citric acid | OA | C6H8O7 | 191,0191 | neg | 1,74 | yes | blackcurrant | blackcurrant, strawberry | [19,38] | [10,57] | [29,30,49,59] | [14,58] |
| Malic acid | OA | C4H6O5 | 133,0133 | neg | 1,47 | yes |  | cherry, strawberry | [19,38] | [7,10,26,57] | [29,30,49,59] | [14,58,60] |
| Quinic acid | OA | C7H12O6 | 191,0555 | neg | 1,33 | yes | cranberry | cherry, cranberry, strawberry | [19,38] | [26] | [29,30,49] | [61] |
| Quinic acid diglycoside | OA | C19H34O17 | 533,1721 | neg | 1,27 | no | cranberry | cranberry | [19,38] |  |  |  |
| Benzoic acid hexoside | BAD | C13H16O7 | 329,0877 | neg | 3,11 | no | cranberry |  |  |  | [29,32,48] |  |
| Hippuric acid | BAD | C9H9NO3 | 180,0655 | pos | 3,37 | no | cranberry |  |  |  | [62] |  |
| Catechin | PC | C15H14O6 | 291,0862 | pos | 2,37 | yes |  |  | [42] | [5,8,10] | [31,46,48] | [36] |
| Epicatechin | PC | C15H14O6 | 291,0862 | pos | 2,77 | yes |  | cherry | [1,42] | [5,10,25] | [31,34,44] |  |
| Procyanidin A2 | PC | C30H24O12 | 577,1352 | pos | 4,55 | yes |  | cranberry |  |  | [63] |  |
| Disaccharide | S | C12H22O11 | 341,1089 | neg | 1,18 | no |  | blackcurrant, strawberry |  |  |  |  |
| hexopolyol | S | C6H12O6 | 181,0708 | pos | 1,22 | no |  | cherry |  |  |  |  |
| Mannitol-hexose | S | C12H24O11 | 343,1245 | pos | 1,2 | no | cherry | cherry |  |  |  |  |
| Carboxylinalool glucoside | T | C16H26O8 | 364,1965 | pos | 3,43 | no | blackcurrant |  |  |  |  |  |
| Linalooloxide glucoside | T | C16H28O7 | 350,2172 | pos | 3,57 | no | blackcurrant |  | [64] |  |  |  |
| Diterpene-hexoside | T | C26H28O9 | 485,1806 | pos | 5,43 | no | blackcurrant |  |  |  |  |  |
| Diterpene2 hexoside | T | C26H30O9 | 487,1962 | pos | 7,1 | no | blackcurrant |  |  |  |  |  |
| Nerolidol hexoside | T | C21H34O6 | 383,2428 | pos | 6,7 | no | strawberry |  |  |  |  |  |
| nerolidol | T | C15H24O | 203,1795 | pos | 7,37 | no | strawberry |  |  |  |  |  |
| Linalyl arabinosylglucoside | T | C21H34O10 | 493,2290 | pos | 6,76 | no | strawberry |  |  |  |  |  |
| Benzoic acid pentosyl hexoside | BAD | C18H24O11 | 434,1656 | pos | 2,37 | no |  |  |  |  | [48] |  |
